# Increased versican and fibrosis in mesenteric lymph nodes disrupts immune surveillance and drives systemic bacterial dissemination in cirrhosis

**DOI:** 10.1101/2025.11.25.690350

**Authors:** Pinky Juneja, Aarti Sharma, Bhaskar Sharma, Guresh Kumar, Deepika Jakhar, Neha Chauhan, Vikas Khillan, Amar Mukund, Dinesh M Tripathi, Shiv K Sarin, Rakhi Maiwall, Savneet Kaur

**Affiliations:** Department of Molecular and Cellular Medicine, Institute of Liver and Biliary Sciences, New Delhi; Department of Hepatology, Institute of Liver and Biliary Sciences, New Delhi; Department of Microbiology, Institute of Liver and Biliary Sciences, New Delhi; Department of Interventional Radiology, Institute of Liver and Biliary Sciences, New Delhi

**Keywords:** Liver cirrhosis, Mesenteric lymph node, Immune Dysfunction, T cell activation, Versican

## Abstract

**Background and Objective:** Mesenteric lymph nodes (MLN) are immunological barriers against bacterial translocation (BT). Enhanced gut BT through MLN facilitates bacterial spread and higher mortality in cirrhosis. We aimed to elucidate mechanisms underlying MLN failure to effectively contain BT during advanced cirrhosis.

**Design:** BT and immune cells were analyzed in lymphoid organs and circulation of control and CCl_4_ models, with and without MLN (MLNx). MLN proteomics identified versican (VCAN) as major upregulated protein in cirrhosis, whose immunomodulatory function was examined *in vitro* and *in vivo* in CCl and (Bile duct ligation) BDL models. Plasma VCAN were measured in end-stage cirrhosis patients and analyzed as mortality predictor.

**Results:** In control rats, bacteria were confined to MLN, whereas cirrhotics showed BT to MLN, lymph, and portal blood. Compared to control, CCl_4_ rats had increased activated Th-cells in MLN but reduced in circulation. In control-MLNx rats, activated Th-cells were reduced in circulation vs controls. In BDL models, MLN CFU correlated with VCAN level*. In vitro*, VCAN enhanced T cell suppression and impaired migration which was reversed by CD44 blockade. *In vivo* VCAN knockdown reduced fibrosis and bacterial burden in MLN, while restoring Th-cell activation locally and systemically. Clinically, plasma VCAN levels were elevated in advanced cirrhosis patients and remained an independent predictor of 28-day sepsis-related mortality.

**Conclusion:** Increased VCAN impairs T cell activation and migration in MLN, fostering immune suppression and bacterial persistence. Plasma VCAN levels serve as promising biomarker for MLN dysfunction and prognostic factor for predicting sepsis-related mortality in end-stage cirrhosis.

**What is already known on this topic** – *Enhanced gut bacterial translocation through mesenteric lymph nodes (MLN) facilitates systemic bacterial spread and increases mortality in cirrhosis. The mechanisms underlying MLN failure to effectively contain bacterial spread during advanced cirrhosis remain largely unknown*.

**What this study adds** – *The study unveils a critical role of lymph node fibrosis and elevated versican (VCAN) expression in causing deranged immune responses and bacterial clearance in MLN, increasing systemic bacterial load and immunosuppression. Most importantly, high plasma VCAN emerges as a prognostic biomarker for functional failure of MLN and 28-day mortality predictor in critically ill patients with cirrhosis*.

**How this study might affect research, practice or policy** – *VCAN, representing enhanced MLN fibrosis and dysfunction, emerges as a predictive biomarker of adverse clinical outcomes in patients with advanced cirrhosis*.

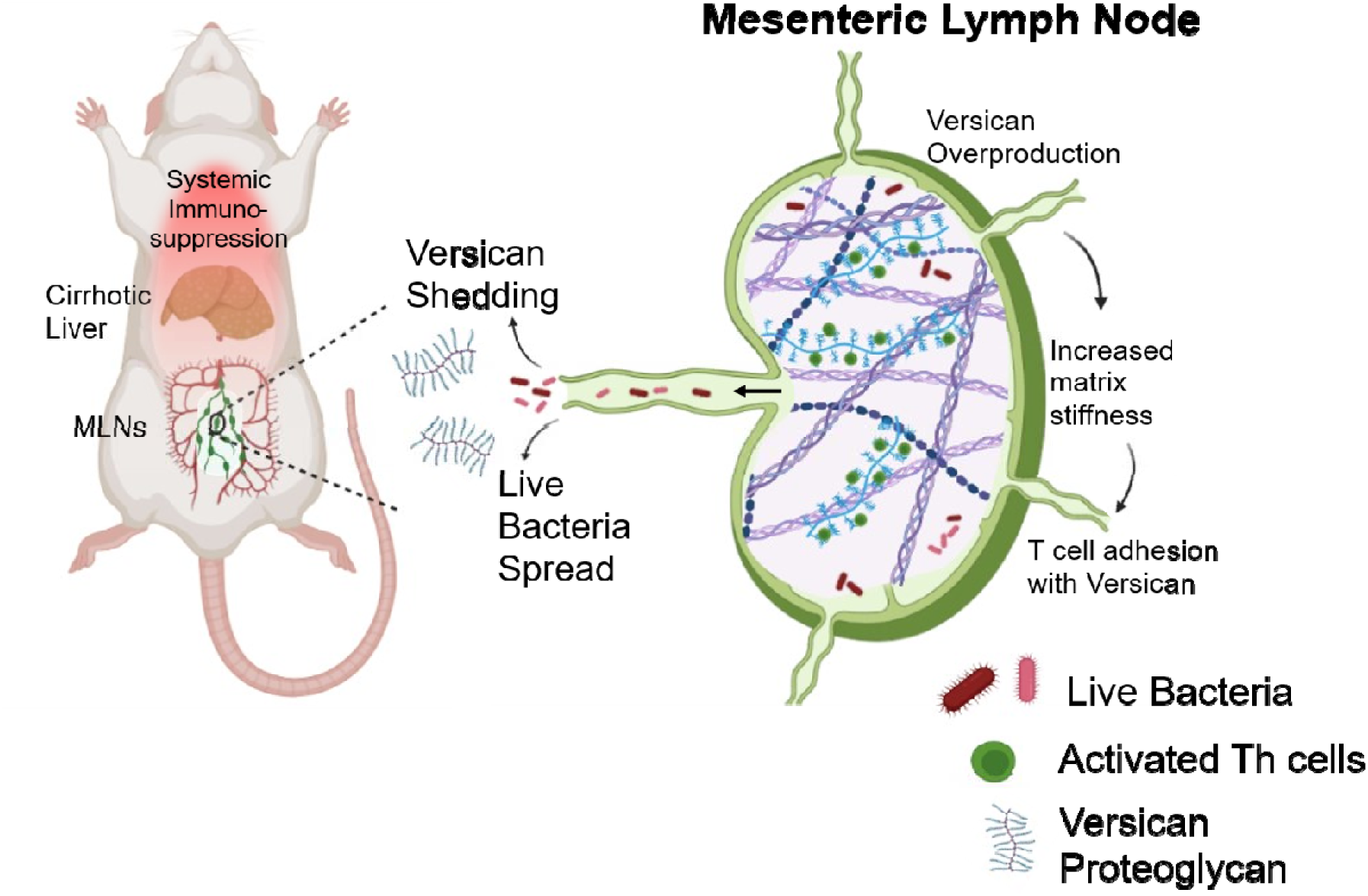

## Introduction

Pathological bacterial translocation (BT), the passage of viable bacteria and/or bacterial products from the gut to mesenteric lymph nodes (MLN) and other extraintestinal sites, is a key driver of disease progression and complications in cirrhosis [1]. BT plays a pivotal role in driving immune dysfunction, infection-related morbidity, and mortality among cirrhosis patients [2]. Under physiological states, MLN serve as the immunological sentinels, playing a central role in intercepting gut-derived microbes and preventing their systemic bacterial dissemination [3]. However, in cirrhosis, factors such as increased intestinal permeability, bacterial overgrowth, and impaired mucosal immunity overwhelm the MLN’s containment capacity, resulting in increased bacterial burden within the MLN [4,5]

Clinical observation in patients with advanced cirrhosis exhibits higher rates of MLN cultures and increased intestinal permeability compared to those with early-stage disease [6]. Supporting this, experimental models using labeled bacteria have demonstrated increased BT from MLN to systemic circulation in rats with advanced cirrhosis [7]. The failure of MLN to effectively contain gut bacteria is a critical event that facilitates bacterial dissemination into systemic circulation, ultimately contributing to complications such as sepsis, spontaneous bacterial peritonitis, hepatorenal syndrome, and hepatic encephalopathy (HE) [8].

Understanding the alterations and mechanisms underlying this failure of MLN is crucial for identifying new therapeutic targets to prevent infection complications and their sequelae in cirrhosis. To address this knowledge gap, we investigated the immune composition and functional capacity of MLN in rat model of CCl -induced cirrhosis. We assessed immune cell profiles and bacterial load within the MLN and systemic, portal, and lymphatic compartments, and evaluated the effect of MLN resection on immunity and BT. Proteomics analysis of MLN identified a marked upregulation of protein versican (VCAN), a chondroitin sulfate proteoglycan, known to influence immune cell behavior in inflammation or fibrosis [9,10]. VCAN has previously been implicated in modulating immune cell migration and response in other organs [11]. However, its role in MLN immune function and BT in cirrhosis has not been explored. We therefore functionally characterized VCAN to determine its contribution to MLN fibrosis, immune cell response, and bacterial containment. Additionally, we extended our findings to a clinical cohort by measuring VCAN levels in plasma and ascitic fluid of advanced cirrhosis patients and analyzing their association with short-term mortality.

## Methods

### Experimental Groups and Animal Model Design

Male Sprague Dawley rats (6 weeks old, 200-250 g) were randomly assigned to experimental groups. Liver fibrosis was induced using intraperitoneal injections of CCl in olive oil (1 mL/kg, twice weekly for 12 weeks). Animals were divided into three main groups: (1) Control (no treatment), (2) C+LPS (control rats receiving 1 mg/kg LPS intraperitoneally), and (3) CCl +LPS (cirrhotic rats receiving a single LPS dose after 12 weeks of CCl). A separate cohort of control rats underwent surgical resection of MLN, followed by division into: (4) CMLNx (MLN-resected rats receiving LPS after 3 weeks), and (5) CCl -MLNx (CCl administration 3 weeks after MLN resection for 12 additional weeks, followed by LPS injection). For VCAN knockdown studies, siRNA against VCAN or scrambled control was delivered in three doses on alternate days via *in vivo* JET-PEI in CCl_4_ rats. BDL models were prepared to study the VCAN level during cirrhosis progression. All data were analyzed using appropriate statistical tests (one-way ANOVA, two-way ANOVA, and Student’s t-test, as applicable), and results were expressed as mean ± SD, with p < 0.05 considered statistically significant.

### Clinical Study Design

This prospective study enrolled 86 critically ill patients over the period of one year with advanced cirrhosis, along with 15 healthy controls (Ethics protocol no. IEC/2024/116/MA05). Patients were not involved in the design and conduct of this research. Patients with prior liver transplantation or malignancy were excluded. Baseline demographic, clinical, and laboratory parameters were recorded at admission, and plasma samples were collected for VCAN quantification. Patients were followed for 28 days or until death, liver transplantation, or TIPS, whichever occurred first. Continuous variables are expressed as median (IQR) and categorical variables as counts (%). Between-group differences were analyzed using the Mann-Whitney U for 2 groups and one-way ANOVA for more than 2 groups. Variables with p < 0.2 in univariate analysis were included in multivariate Cox regression to identify independent predictors of 28-day mortality, reported as hazard ratios (HRs) with 95% CIs. The prognostic performance of VCAN was evaluated using ROC curve analysis (AUC, 95% CI) and the Youden index to determine the optimal cutoff. Kaplan-Meier curves with log-rank tests were used for survival comparisons. Because liver transplantation and TIPS preclude the primary event (death), competing-risk regression was performed using the Fine and Gray model to account for these outcomes.

## Results

### MLN as a critical barrier against bacterial dissemination in liver cirrhosis

In liver cirrhosis patients, pathological gut BT leads to complications and increased mortality. To study the contribution of MLN in BT to other extraintestinal sites, we first developed animal models of liver cirrhosis using CCl_4_ for 12 weeks and quantified indigenous bacterial load in different organs of cirrhosis and control SD rats (**Supp Fig. 1a, b**). Control rats showed no detectable bacterial colonies in MLN or extraintestinal sites, whereas cirrhotic rats harbored significant bacterial loads in MLN of 80% of cirrhosis rats, which also showed ascites (**Supp Fig. 1c, d**). Bacterial load was also present in the lung, liver, blood, and lymph of CCl_4_ rats. To study BT in cirrhosis, we administered GFP-labeled bacteria orally and quantified the CFU (**Fig. 1A**). In healthy rats, bacteria localized exclusively to the MLN (**Fig. 1B**). In contrast, cirrhotic rats presented significantly higher bacterial count in MLN, and dissemination to extraintestinal organs (**Fig. 1B-D**), including efferent mesenteric lymph and portal blood.

**Figure 1:**
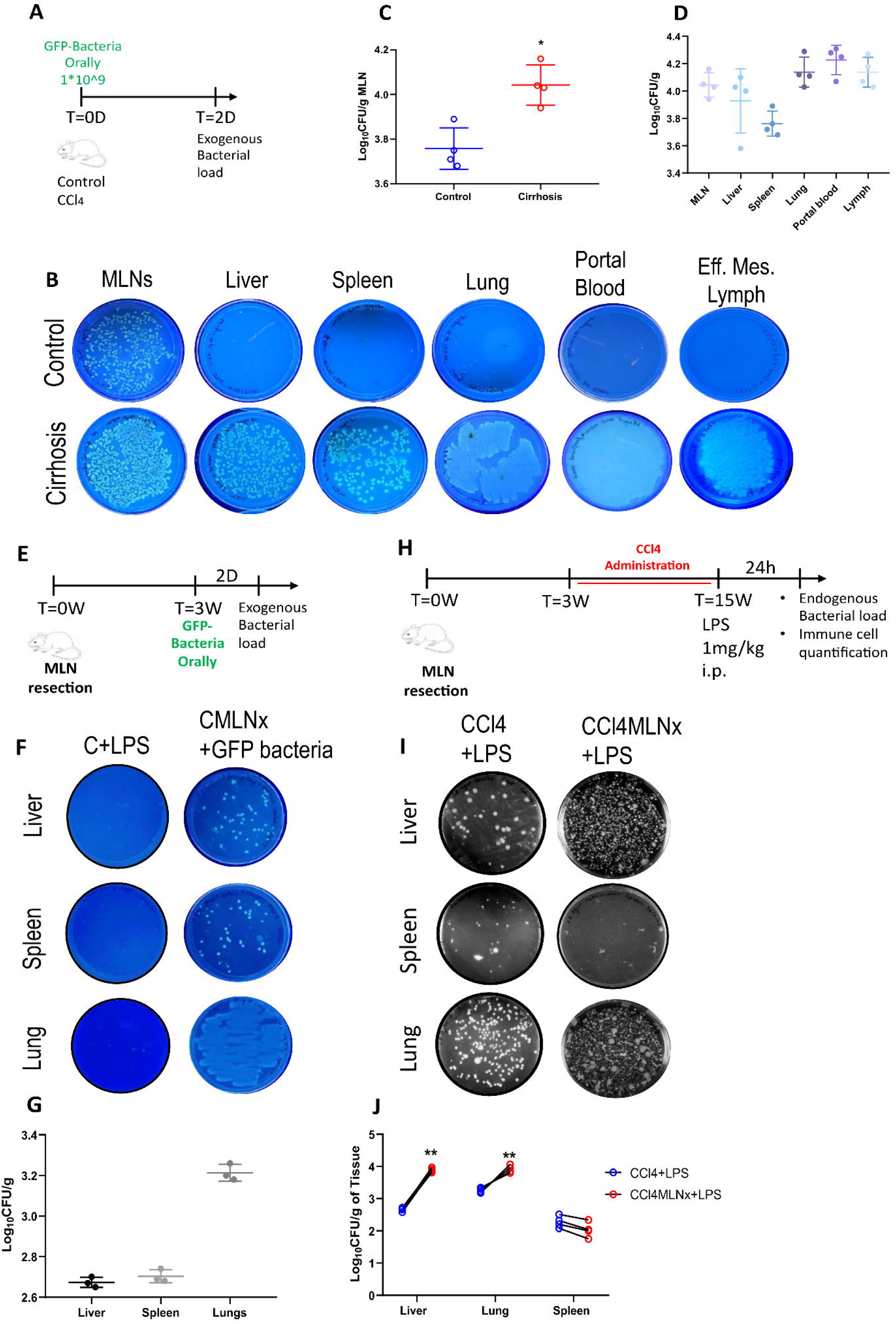
Increased bacterial load and bacterial translocation from MLN to other extraintestinal organs: (A) Schematic of experimental design showing oral gavage of 10^9^ GFP-labeled bacteria followed by sample collection after 48 h. (B) Representative culture plates of MLN, liver, spleen, lung, portal blood, and efferent mesenteric lymph from control cirrhotic rats. (C) Quantification of CFU in MLN of control and cirrhotic rats. (D) Quantification of CFU in MLN, liver, spleen, lung, portal blood, and efferent mesenteric lymph of cirrhotic rats. (E) Schematic of experiment design showing MLN resection (CMLNx) followed by oral administration of 10^9^ GFP-labeled bacteria in control rats. Samples were collected after 48h. (F–G) Detection and quantification of GFP-bacteria in the liver, spleen, and lungs of control rats with and without MLN. (H) Experimental timeline for induction of cirrhosis in MLN-resected rats (CCl -MLNx). (I–J) Detection and quantification of endogenous bacterial load in liver, spleen, and lungs of CCl and CCl -MLNx rats. Data were polled from >3 independent experiments and expressed as mean+SD. Statistics assessed by (C) unpaired two-tailed Student’s t-test and (J) Two-way ANOVA with Tukey’s multiple comparison test. *p< 0.05, **p< 0.01, ***p< 0.001. MLN: Mesenteric Lymph Nodes, CFU: Colony-forming unit.

To delineate the role of MLN in containing bacterial infection, MLN were surgically removed from control rats (CMLNx), and bacterial loads in various organs were measured 24 h after LPS challenge (**Supp Fig. 1e**). Endogenous bacterial load was observed in lungs of CMLNx **(Supp Fig. 1f),** indicating MLN are essential to limit BT beyond the gut. However, exogenous GFP-labeled bacterial load was present in the liver, spleen, and lungs of CMLNx rats, indicating the indispensable role of MLN in preventing BT (**Fig. 1E-G**). We observed no change in liver architecture in CMLNx vs control. To assess the role of MLN in cirrhosis progression, some MLN-resected control rats were administered CCl for 12 weeks (CCl -MLNx) (**Fig. 1H**). In CCl_4_-MLNx rats, bacterial loads were significantly increased in the lungs and liver compared to CCl_4_ rats (**Fig. 1I, J**), accompanied by elevated plasma endotoxin levels (**Supp Fig. 1g**). Intriguingly, MLN removal accelerated liver fibrosis progression, collagen deposition, and decreased survival compared to CCl rats with intact MLN (**Supp Fig. 1h-j**). Before LPS administration, up to 12 weeks of the CCl model, there was a single (7%) mortality in cirrhotic rats with intact MLNs as compared to 4(30%) in MLN resected cirrhotic rats. After LPS administration, mortality increased to 4(30%) in CCl_4_-MLNx rats compared to 2(15%) in CCl_4_ rats with intact MLNs **(Supp Fig. 1h).** These findings establish that healthy MLN act as a crucial barrier preventing bacterial dissemination beyond the gut. In cirrhosis, MLN dysfunction exacerbates BT and its systemic spread, underscoring the importance of MLN integrity in immune defense failure associated with cirrhosis.

### Activated CD4 T cells are increased in MLN but reduced in the blood of cirrhotic animals

Under physiological conditions, bacterial components are transported to the MLN by dendritic cells (DC), where they activate macrophages and T cells to initiate immune responses. To delineate compartment-specific immune alterations, we analyzed MLN, peripheral blood, portal blood, and efferent mesenteric lymph from control (C+LPS) and cirrhotic (CCl4+LPS) rats following a single LPS challenge (1 mg/kg). The gating strategy is shown in **Supp Fig. 2**. In MLN, CD103, CD80 antigen-presenting DCs, and CD68 macrophages were markedly increased in CCl +LPS rats, whereas inflammatory monocytes decreased and non-inflammatory monocytes increased relative to C+LPS (**Fig. 2A–C**). CD4 and CD8 T cell frequencies remained comparable between groups (**Supp. Fig. 3a**). However, CD4 CD134 activated T cells were significantly elevated in both C+LPS and CCl +LPS groups vs control; an additional rise in CD8 CD25 Tregs was observed only in cirrhotic MLN (**Fig. 2D**). Correspondingly, TGF-β levels were increased, and IL-10 and TNF-α levels were reduced while IL-6 remained unchanged in cirrhotic MLN, suggesting an attenuated proinflammatory activity and a shift toward an immunosuppressive milieu (**Fig. 2E**).

**Figure 2.**
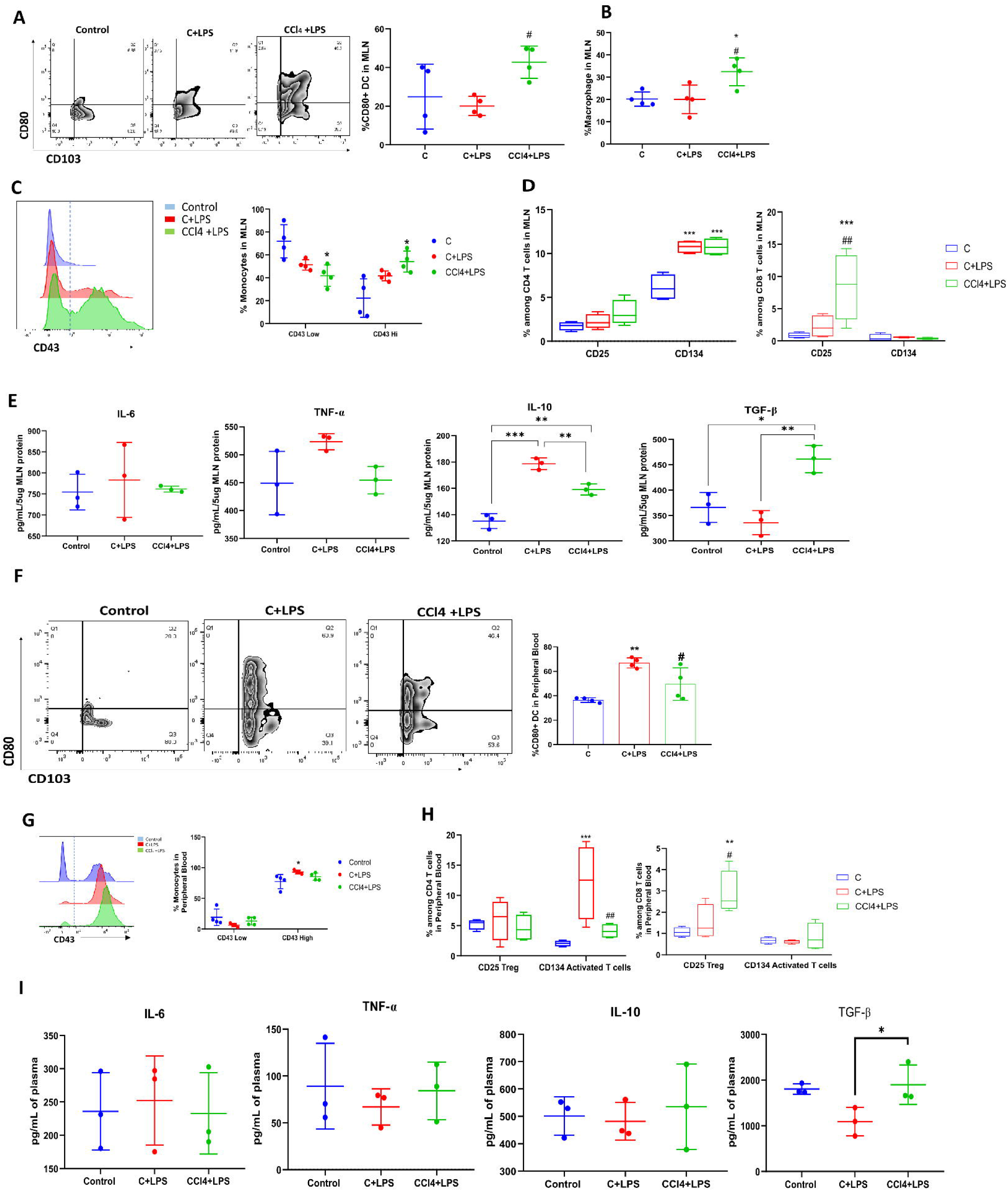
Suppressed immune response in the MLN and peripheral blood of cirrhotic rats. (A) Representative plots and quantification of CD80 CD103 DC gated on CD45 CD11b cells in MLN of control, C+LPS, and CCl +LPS rats. (B) Quantification of CD68 macrophages in MLN (C) Expression and quantification of CD43 hi and low monocytes frequency gated on CD45 CD11b cells in MLN. (D) CD134 activated and CD25 Tregs among the CD4^+^ and CD8^+^ T cell subsets in MLN. (E) Cytokine level in the MLN lysate. (F) Representative plots and quantification of CD80 CD103 DC in circulation. (G) Expression and quantification of CD43 hi and low monocytes frequency gated on CD45 CD11b/c cells in the peripheral blood of the respective group. (H) Proportions of CD134 activated and CD25 Tregs within CD4 and CD8 subsets in peripheral blood. (I) Cytokine levels in the systemic circulation. (A-I) Data pooled from >3 independent experiments and presented as mean+SD. Statistics assessed by (A, B, E, F, and I) One-way ANOVA and (C, D, G, and H) Two-way ANOVA with Tukey’s multiple comparison. *p< 0.05, **p< 0.01, ***p< 0.001. *vs control; #vs C+LPS. MLN: Mesenteric Lymph Nodes; DC: Dendritic cells.

In the systemic circulation, DC frequency was reduced in CCl +LPS compared with C+LPS (**Fig. 2F**). Overall, monocyte, CD4, and CD8 T cell fractions were unchanged (**Fig. 2G**, **Supp Fig. 3b**). Notably, CD4 CD134 activated T cells were significantly decreased, while CD8 CD25 Tregs were increased in CCl +LPS vs C+LPS rats (**Fig. 2H**). Cytokine profiling revealed no change in IL-6, IL-10, or TNF-α levels but showed elevated TGF-β, consistent with a systemic immunosuppressive profile (**Fig. ╁2I**).

In the portal circulation, DC frequency was increased in the CCl +LPS group compared with C+LPS. This was accompanied by an increase in inflammatory and a reduction in non-inflammatory monocytes **(Supp Fig. 3c, d).** CD4 and CD8 T cell proportions were unaltered (**Supp Fig. 3e**). However, CD4 CD134 activated T cells were decreased, whereas CD8 CD25 Tregs were significantly elevated in the portal blood of CCl +LPS rats compared with C+LPS **(Supp Fig. 2f).** In the efferent mesenteric lymph, CD4 and CD8 T cell frequencies remained unchanged between groups **(Supp Fig. 2g).** Notably, CD134 expression on both CD4 and CD8 T cells was significantly reduced in the CCl +LPS group, with no corresponding change in Tregs (**Supp Fig. 2h**). In summary, cirrhotic rats exhibited distinct compartmentalized immune alterations characterized by increased immune activation and antigen-presenting activity in the MLN with ongoing systemic immunosuppression, reflecting impaired T cell responsiveness and a shift toward a regulatory, tolerance-prone phenotype in the blood that may contribute to immune dysfunction in advanced liver disease.

### MLN resection reduces activated CD4 T cells in systemic circulation

To assess the contribution of the MLN to systemic immune changes in cirrhosis, immune cell profiles and cytokine responses were compared in peripheral and portal compartments following MLN resection. In healthy rats, MLN resection (CMLNx+LPS) led to reduced DC frequency, increased inflammatory and decreased non-inflammatory monocytes in systemic circulation compared with C+LPS rats (**Fig. 3A, B**). The CD4 T cell fraction increased while CD8 T cells decreased, resulting in a higher CD4/CD8 ratio (**Supp Fig. 4a**). Critically, the proportion of CD134 activated CD4 T cells was markedly reduced in CMLNx rats, without changes in Tregs (**Fig. 3C**). There were no significant alterations in circulating IL-6, IL-10, TNF-α, or TGF-β levels in CMLNx vs C+LPS (**Fig. 3D**). In the portal circulation of CMLNx rats, DC frequency remained unchanged while inflammatory monocytes decreased compared to C+LPS rats (**Supp Fig 4b, c**). The portal CD4 T cell fraction increased alongside reduced CD8 and CD4 CD134 T cells (**Supp Fig 4d, e**).

**Figure 3:**
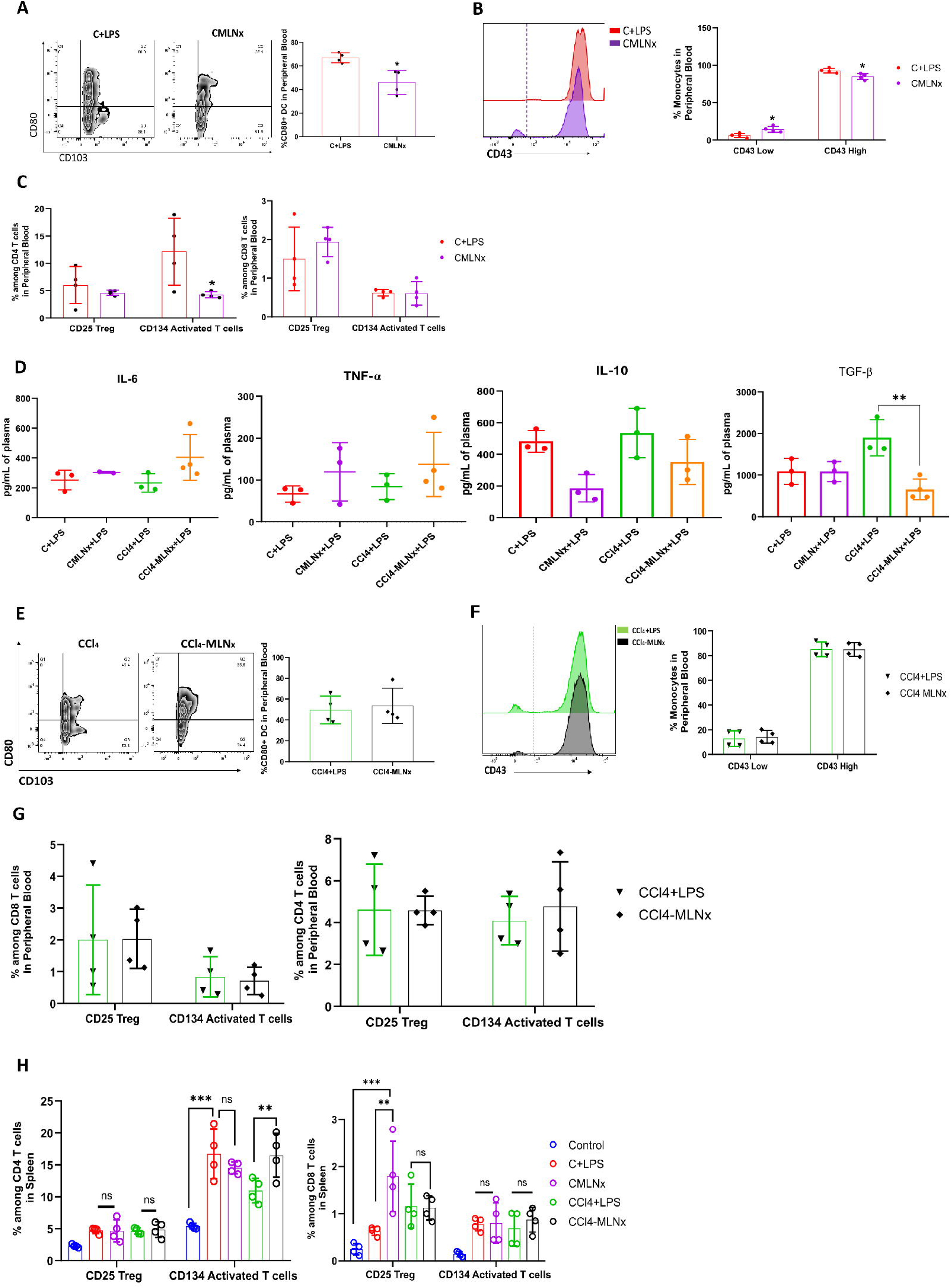
MLN resection reduced activated T cells in the peripheral blood of controls but not in CCl□ rats. (A) Representative plots and quantification of CD80 and CD103 DC in the peripheral blood of control rats with/without MLN. (B) Expression and quantification of CD43 hi and low monocyte frequency in peripheral blood. (C) Quantification of CD134 activated T cells and CD25 Tregs among the CD4 and CD8 T cell subsets in the peripheral blood of control and CMLNx rats. (D) Cytokine level in systemic circulation. (E) Representative plots and quantification of DC in the peripheral blood of cirrhotic rats with/without MLN. (F-G) Expression and quantification of (F) CD43 hi/low monocyte frequency and (G) CD134 activated T cells and CD25 Tregs among the CD4 and CD8 T cell subsets in peripheral blood of cirrhotic rats with and without MLN. and (H) Quantification of CD134 activated T cells and CD25 Tregs among the CD4 and CD8 T cell subsets in the spleen of respective groups. (A-K) Data pooled from >3 independent experiments and presented as mean+SD. Statistics assessed by (A, D and E) Unpaired two-tailed Student’s *t*-test and (D, C, F, G, and H) Two-way ANOVA with post-hoc Tukey’s multiple comparison test. *p< 0.05, **p< 0.01, *** p< 0.001. MLN: Mesenteric Lymph Nodes; DC: Dendritic cells.

In cirrhotic rats, MLN resection did not affect DC, monocytes, or T cell subset frequencies, though the CD4/CD8 ratio was higher in CCL4-MLNx rats (**Fig. 3E-G**, **Supp Fig. 4f**). Cytokine analysis showed no significant differences except for decreased TGF-β in CCl4-MLNx relative to CCl4 (**Fig. 3D**). Portal blood DC were unchanged, but non-inflammatory monocytes and CD4/CD8 ratio were elevated in CCl -MLNx rats as compared to CCl4 rats with intact MLN (**Supp Fig.4g-i**). CD134 activated T cell or CD25 Tregs showed no significant change (**Supp Fig. 4j**).

To assess whether the spleen, being another secondary lymphoid organ, compensates for MLN function after resection, we analyzed splenic immune composition across all experimental groups. The frequency of DC and monocytes remained unchanged among groups (**Supp Fig. 4k, l**). CD68 macrophages were reduced in CCl -MLNx rats compared with CCl (**Supp Fig. 4m**). Among T cell subsets, CD4 and CD8 frequencies were comparable; however, the CD4/CD8 ratio increased in CMLNx but remained unchanged in CCl -MLNx (**Supp Fig. 4n**). CD134 activated CD4 T cells decreased in CMLNx rats relative to C+LPS but increased in CCl -MLNx compared with CCl (**Fig. 3H**). Conversely, CD8 Tregs were significantly elevated in CMLNx vs. C+LPS but remained unchanged in cirrhotic groups. Splenic TNF-α mRNA expression decreased following MLN resection in healthy rats, while IL-6, IL-10, and TGF-β transcripts were unchanged (**Supp Fig. □4o**). Overall, MLN resection led to minor changes in splenic immune composition, characterized by altered CD4 T cell activation in healthy rats but minimal compensatory response in cirrhotic conditions. To conclude, MLN resection in healthy rats caused a significant decrease in circulating activated CD4 T cells and modestly altered splenic immune balance, whereas its effects were minimal in cirrhotic rats, likely due to baseline MLN dysfunction.

### Versican upregulation impairs T cell activation and migration in cirrhotic MLN

MLN from control, C+LPS, and CCl_4_+LPS rats were subjected to proteomic analysis to elucidate the molecular mechanism underlying MLN dysfunction in cirrhosis. Comparative quantification of 2220 proteins identified 261 with significant differential expression (fold change ≥1.5 and *P_adj_*<0.2) (**Supp Fig 5a**). Compared to control rats’ MLN, 69 proteins were significantly upregulated and 59 downregulated in the MLN of C+LPS rats, while 126 proteins were upregulated and 92 downregulated in CCl +LPS rats’ MLN. In C+LPS vs CCl_4_+LPS MLN, 47 proteins were significantly altered. Of these, 20 proteins were upregulated, and 27 proteins were downregulated in CCl_4_ (**Fig. 4A**). The heat map shows the differential expression of the top 20 significant proteins between the groups with a fold change of more than 1.5 (**Fig. 4B**). The upset plot reveals that the number of differentially expressed proteins (DEP) is significantly higher in the CCl_4_+LPS group (111 DEP) compared to the C+LPS group (25 DEP) relative to the control (**Fig. 4C**, **Supp Table 1**). In cirrhotic MLN, the enrichment of biological processes related to downregulated proteins revealed the suppression of pathways associated with cellular responses to LPS and bacterial stimuli (**Fig. 4D**). Versican (VCAN), an extracellular matrix (ECM) protein, was especially increased (fold change: 425.6) and originated predominantly from fibroblastic reticular cells (FRCs) within MLN, with expression significantly higher in cirrhotic FRCs relative to controls. Conversely, cirrhotic MLN showed upregulation of proteins related to defense response to bacterial and chemotaxis, such as granzyme, CCR10, and UNC13B, compared to C+LPS. Enrichment of pathways related to significantly upregulated proteins (VCAN, SRGAP2, PTPN9, and DPYSL5) highlights negative regulation of cell migration, cytoskeletal severing, and proliferation (**Supp Fig 5b**).

**Figure 4.**
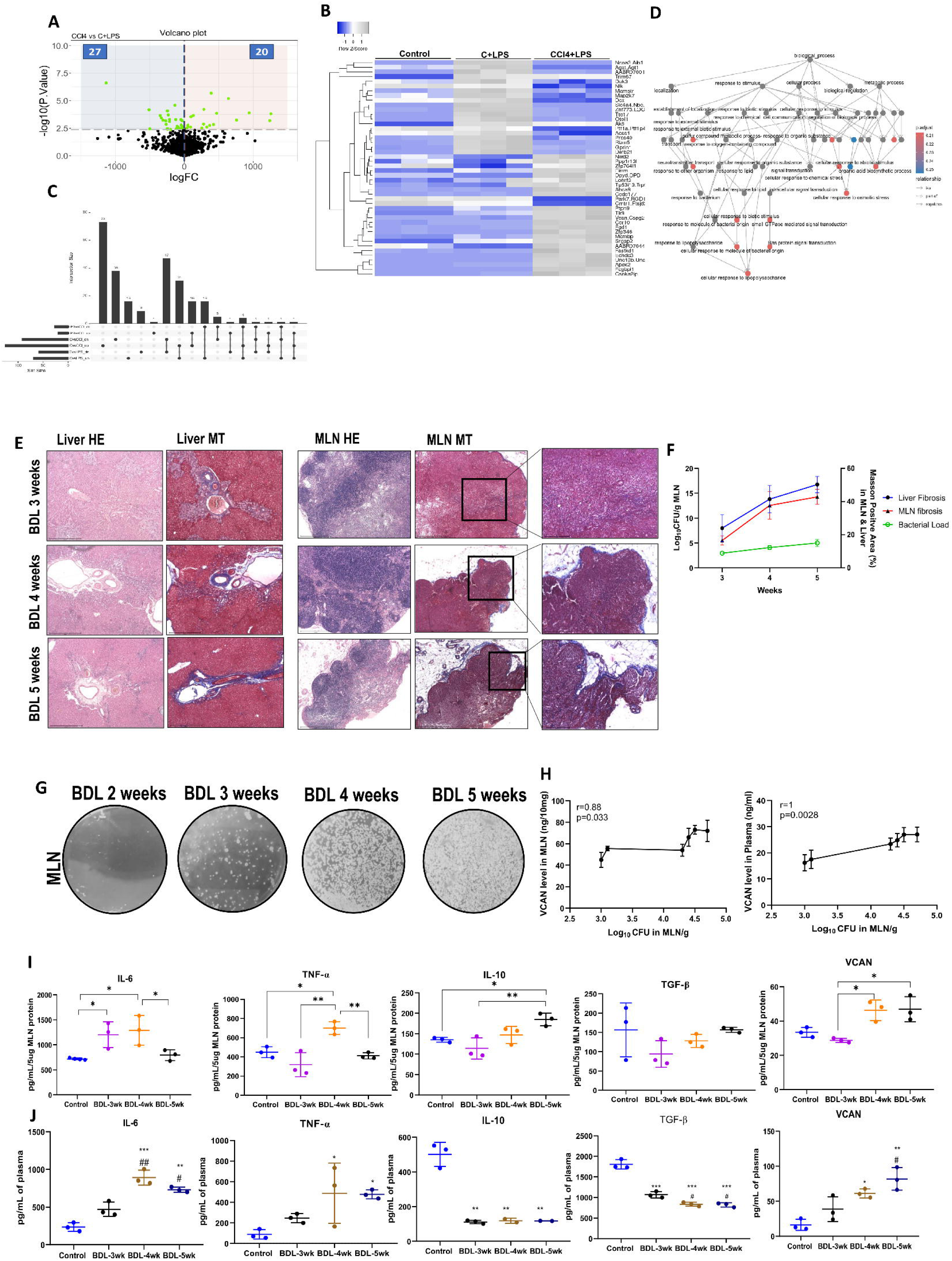
Proteomic profiling and functional role of VCAN in immune regulation. (A) A volcano plot shows differentially expressed proteins (DEP) between C+LPS and CCl +LPS MLN, highlighting proteins with fold changes≥1.5 and P_adj_ < 0.2. (B) A heatmap displays the top 20 significantly altered proteins across groups, with hierarchical clustering. (C) An upset plot summarizes the number and overlap of DEPs among the control, C+LPS, and CCl +LPS MLN. (D) Gene ontology enrichment analysis reveals biological processes associated with significantly downregulated proteins in CCl +LPS MLN, including cellular response to LPS and bacterial stimuli. (E) HE and MT staining of Liver and MLN from BDL models sacrificed at 3, 4, and 5 weeks post-surgery. (F) Graph represents CFU in MLN on the left Y axis and % area of MT staining (fibrosis) on the right Y axis in BDL rats at 3, 4, and 5 weeks. One data point in graph represents the mean of >3 independent experiments. (G) Representative culture plates of MLN showing endogenous bacterial load at different time points in the BDL model. (H) Correlation of bacterial load in MLN with VCAN level in MLN and plasma of BDL model at different time points (n=3 each). (I) (I, J) Quantification of cytokines from (I) MLN Lysate and (J) plasma of BDL models at different time points (n=3 each). (E-J) Data pooled from 3 independent experiments and expressed as mean ± SD. Statistics assessed by (H) Spearman’s rank correlation test, (I, J) One-way ANOVA with post-hoc Tukey’s multiple comparison test. *p < 0.05, **p < 0.01, and ***p < 0.001. * vs control; # vs BDL-3wk. MLN: Mesenteric Lymph Nodes; BDL: Bile duct ligation.

In view of the fact that, as compared to control MLN, we observed an increased percentage of activated CD4 T cells in cirrhotic MLN but not in systemic circulation, we focused on the proteins involved in cell migration. In this regard, one of the proteins, VCAN (fold change: 425.6), was identified as a highly upregulated protein in cirrhotic MLN. Therefore, we further studied the dynamics of VCAN expression during cirrhosis progression. For this, we used a BDL model and analyzed VCAN levels and bacterial load as the disease progressed (**Fig. 4E**). Disease progression was accompanied by increasing fibrosis in both the liver and MLN, along with increased bacterial load in MLN (**Fig. 4F**, **Supp Fig. 5c).** All BDL rats showed the presence of ascites at week 5. Endogenous bacterial load in MLN was first detected in some rats at week 3 post-BDL and continued to rise thereafter (**Fig. 4G**). VCAN levels were significantly higher in MLN of BDL rats with BT as compared to those without BT, irrespective of the presence of ascites **(Supp Fig. 5d).** MLN CFU counts positively correlated with VCAN levels in both MLN and plasma across time points (**Fig. 4H**). MLN cytokine profile showed progressive increases in IL-10 and TGF-β, while IL-6 and TNF-α peaked transiently (**Fig. 4I**). In plasma, IL-10 and TGF-β levels remained stable, whereas IL-6 and TNF-α exhibited a similar transient peak (**Fig. 4J**). VCAN levels increased as the disease progressed in both MLN and plasma. These findings suggest that VCAN expression in the MLN rises with cirrhosis progression, paralleling increased fibrosis and BT.

### VCAN silencing enhances systemic T cell activation and reduces bacterial dissemination

Next, to identify the cellular source of VCAN, cells from control and cirrhotic rat MLN were sorted into immune, stromal fibroblastic reticular cell (FRC), and endothelial populations using specific markers, followed by quantification of VCAN expression in each subset (**Supp Fig. 6a**). Among sorted cell populations, FRC had the highest VCAN expression, with significantly greater relative expression in cirrhotic FRC compared to control FRC (**Supp Fig. 6b, c**). Given VCAN’s role in T cell activation and migration, we further investigated its functional relevance in the *in vitro* assays using recombinant VCAN (rVCAN) protein. We isolated and cultured FRCs from control MLN and characterized them using the PDPN marker (**Fig. 5A**). FRCs in the presence of rVCAN resulted in T cell aggregation compared to when cultured without rVCAN (**Fig. 5B-C**). To determine if VCAN affects the activation of T cells, FRCs were exposed to LPS with or without rVCAN. Sorted CD3 T cells from control rats’ MLN were co-cultured for 6 hrs, and flow cytometry analysis revealed a slight decrease in % CD4 CD134 activated T cells and a significant increase in % of CD8 CD25 Treg cells in the presence of rVCAN (**Fig. 5D**). Transwell migration assay demonstrated a reduced T cell migration from the upper to lower chamber when FRC was cultured with rVCAN compared to those without rVCAN protein (**Fig. 5E**). To determine whether VCAN-mediated effects on T cells were dependent on CD44 signaling, as suggested previously [12], we blocked the CD44 receptor on T cells using blocking antibody. This resulted in restored T cell migratory responses, indicating that the FRC-VCAN-CD44 interaction acts as a critical regulatory axis (**Fig. 5E**). To functionally evaluate the role of VCAN in T cell suppression, we co-cultured CD134 activated T cells with CD25 Tregs in the presence or absence of rVCAN (**Fig. 5F**). rVCAN enhanced Treg-mediated suppression compared to Tregs alone, as indicated by decreased expression of CD134 marker on T cells and reduced TNF-α secretion (**Fig. 5F, G**). These results suggest that VCAN interacts with CD44 on T cells, suppresses migration and contributes to an immunosuppressive microenvironment in MLN.

**Figure 5.**
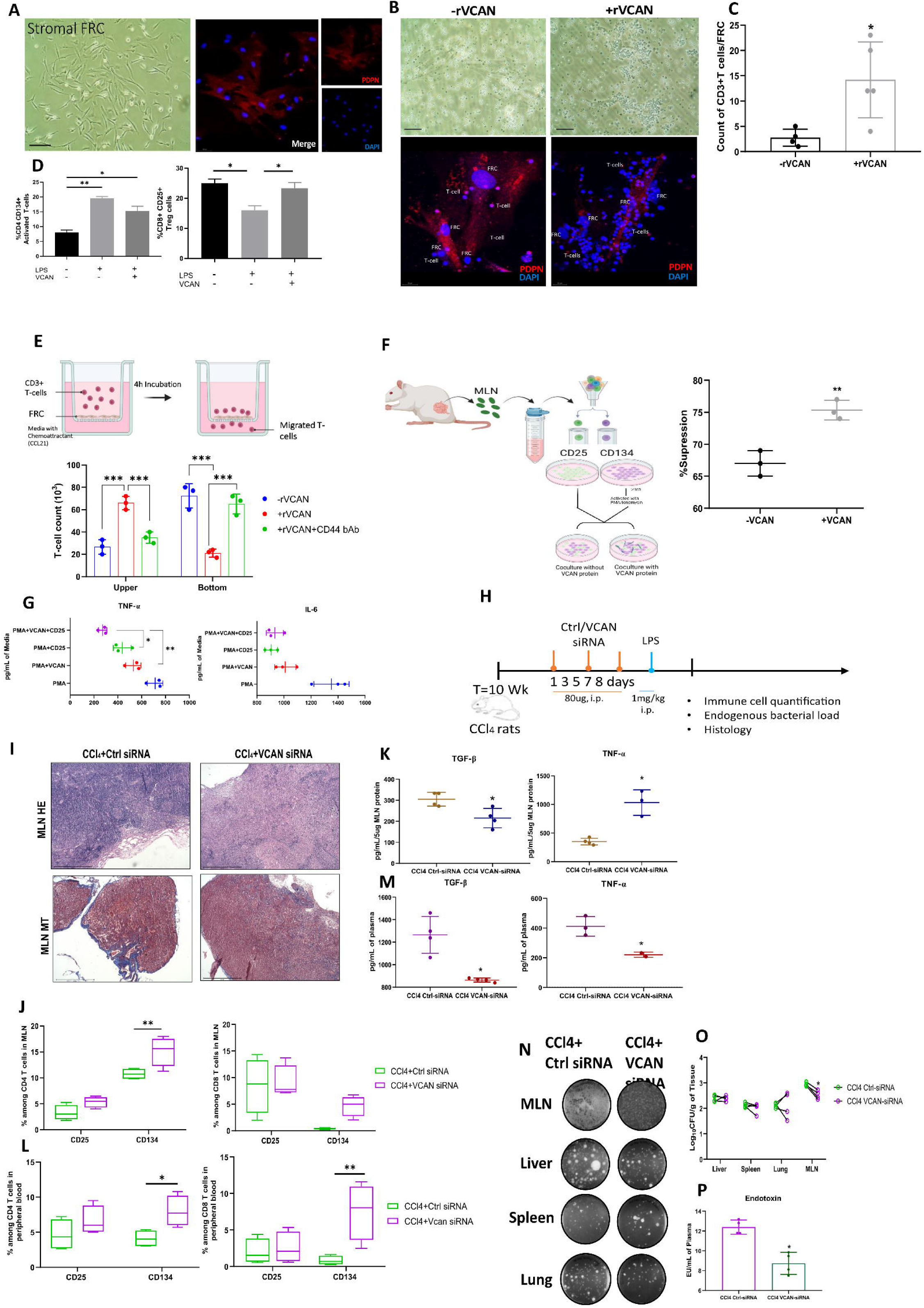
*In vivo* knockdown of VCAN increased activated T cell response in peripheral blood. (A) Representative image of *in vitro* culture of FRCs isolated from MLN and imaging with PDPN marker. (B) *In vitro* co-culture of FRC with T cells in the +/- of recombinant VCAN (rVCAN) protein. Brightfield and IF images show T cell aggregation on FRCs cultured with rVCAN. (C) Quantification of T cells per FRC. (D) Quantification of CD134 activated T cells and CD25 Tregs following co-culture of FRCs with CD3 T cells in the +/- of rVCAN and LPS. (E) Schematic of transwell migration assay of T cells in response to FRCs cultured with or without rVCAN and further quantification of T cells from the top and bottom chambers of transwell. (F) Schematic of T cell suppression assay and further quantification of % suppression of activated CD134 T cells in the +/- of rVCAN. (G) Cytokine quantification of media collected from different conditions. (H) Schematic of the siRNA treatment timeline. (I) HE and MT staining of control and VCAN siRNA-treated MLN. (J) Quantification of activated T cells and Treg cells among the T cell subsets in MLN of VCAN and ctrl siRNA-treated cirrhotic rats. (K) Cytokine and VCAN levels in MLN lysates of the studied groups (n > 3). (L) Quantification of activated T cells and Treg cells in the peripheral blood of VCAN and ctrl siRNA-treated cirrhotic rats. (M) Cytokine quantification in plasma of the studied groups (n>3). (N-O) Quantification of endogenous bacterial load in 100mg of MLN, liver, spleen, and lung of VCAN and ctrl siRNA-treated cirrhotic rats (n=3 each). (P) Plasma endotoxin quantification from control and VCAN siRNA-treated cirrhotic rats (n=4 each). Data pooled from >3 independent experiments and expressed as mean ± SD. Statistics assessed by (C, F, K, M, P) Unpaired two-tailed Student’s *t*-test, (D, G) One-way ANOVA (E, J, L, O) Two-way ANOVA with post-hoc Tukey’s multiple comparison test. *p < 0.05, **p < 0.01, and ***p < 0.001. * vs control. FRC: Fibroblastic reticular cells, PDPN: Podoplanin, MLN: Mesenteric Lymph Nodes; rVCAN: recombinant Versican protein.

Next, to define VCAN’s role *in vivo*, we performed VCAN knockdown in cirrhotic rats (**Fig. 5H**). *In vivo,* VCAN-specific siRNA duplexes versus control (ctrl) siRNA duplexes were used with Polyplus *in vivo*-jetPEI as the transfection medium and injected i.p. qPCR confirmed efficient VCAN silencing with ∼90% reduction in VCAN mRNA expression in MLN of siRNA-treated rats compared to ctrl-siRNA-treated rats (**Supp Fig. 6d**). After confirming VCAN inhibition *in vivo*, a single dose of LPS (1 mg/kg) was injected to study subsequent immune responses and bacterial load in MLN and systemic circulation. Histological analysis using MT staining revealed a reduction in MLN fibrosis in VCAN-knockdown rats with no such effect on the liver (**Fig. 5I**, **Supp Fig. 6e**). Flow cytometry analysis of MLN showed a significantly increased proportion of DC, inflammatory monocytes, and CD4 activated T cells in VCAN siRNA-treated CCl4 rats compared to ctrl siRNA-treated CCl_4_ rats (**Fig. 5J**, **Supp Fig. 6f**). Immunosuppressive cytokines (IL-10, TGF-β) and VCAN levels decreased, and TNF-α was significantly increased in MLN after VCAN silencing, while IL-6 remained unchanged (**Fig. 5K**, **Supp Fig. 6g**). Systemically, VCAN silencing enhanced CD134 activated CD4 and CD8 T cells and DCs in blood (**Fig. 5L**, **Supp Fig. 6h**), without affecting other immune cell subsets or splenic composition (**Supp Fig. 6i**). Cytokine profile revealed a significant decrease in TNF-α, TGF-β, and VCAN levels in plasma of the VCAN siRNA group compared to the ctrl-siRNA group (**Fig. 5M**, **Supp Fig 6j**).

The endogenous bacterial loads in MLN and endotoxemia were significantly reduced in VCAN siRNA-treated cirrhotic rats compared to ctrl-siRNA-treated rats; however, in lungs, spleen, and liver, bacterial load was comparable between VCAN and ctrl-siRNA-treated rats (**Fig. 5N-P**). Collectively, these results indicate an immunosuppressive role of VCAN in cirrhotic MLN, which, when silenced, restores effective T cell activation and reduces MLN bacterial burden.

### Circulating VCAN levels are elevated in cirrhosis and independently predict 28-day mortality

We next evaluated circulating VCAN levels in patients with cirrhosis. A prospective study was conducted in 86 hospitalized patients admitted to the ICU and HDU with decompensated end-stage cirrhosis to assess the prognostic value of plasma VCAN as a marker of MLN dysfunction (**Fig. 6A**). Fifteen healthy controls were included for comparison. Baseline plasma VCAN levels were significantly higher in cirrhotic patients (16.8 ng/mL) compared with controls (9.74 ng/mL, p < 0.0001; **Fig. 6B**, **Table 1**). In the cirrhosis group, the majority of patients were male (90%), with alcohol-related liver disease (62.7%) as the predominant etiology. The median MELD-Na score was 29 (range 11-53). Common complications included ascites (88.3%), HE (76.7%), and acute kidney injury (AKI, 75.6%). Bacterial infections were detected in 58 (67.4%) patients.

**Figure 6.**
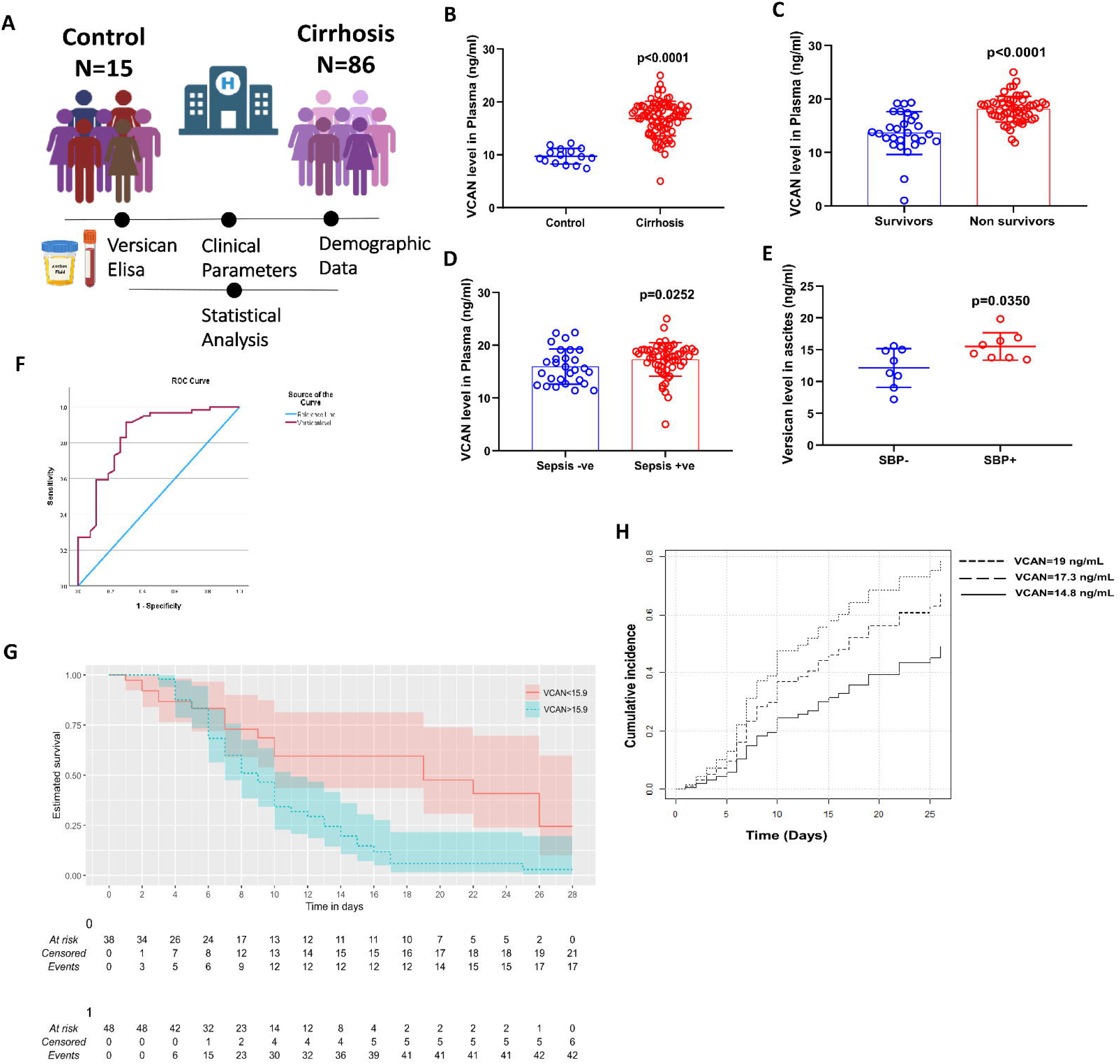
Plasma VCAN levels as a predictor of 3-month mortality in advanced cirrhosis patients. (A) Schematic overview of the prospective study design assessing the prognostic utility of plasma VCAN in patients with advanced cirrhosis. (B) Quantification of plasma VCAN levels in patients with cirrhosis (n = 86) compared to healthy controls (n = 15, p<0.0001). (C) Quantification of plasma VCAN levels among cirrhosis patients between non-survivors (n = 59) and survivors (n = 27; p<0.0001). (D) Quantification of plasma VCAN levels in cirrhosis patients with sepsis (n = 58) vs non-sepsis (n = 28, p = 0.0252). (E) Quantification of plasma VCAN levels in ascitic fluid of patients with/without SBP (p = 0.035). (F) ROC curve for VCAN with AUC = 0.841, 95% CI: 0.744-0.937, p < 0.0001). (G) Kaplan-Meier survival analysis showed a lower survival rate in the high VCAN group (>15.9 ng/mL) than in the low VCAN group (χ² = 9.494, df = 1, p = 0.002). (H) Cumulative incidence functions estimated using the Fine and Gray SHR model to account for competing events (transplantation or TIPS). Solid, dashed, and dotted lines represent VCAN quantiles (25^th^=14.8, 50^th^=17.3, 75^th^=19 ng/mL). (B-E) Data expressed as mean ± SD. Statistical comparisons were made using the Mann-Whitney U test. (G-H) Data are presented as median survival (95% CI). Statistical comparisons were made using the log-rank test.

**Table 1:**
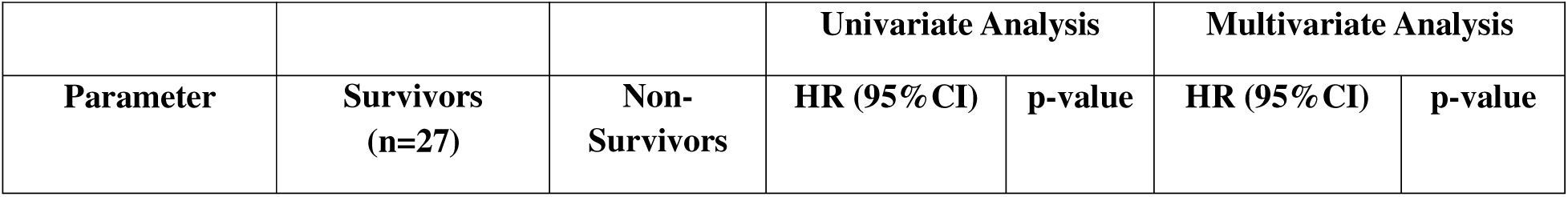
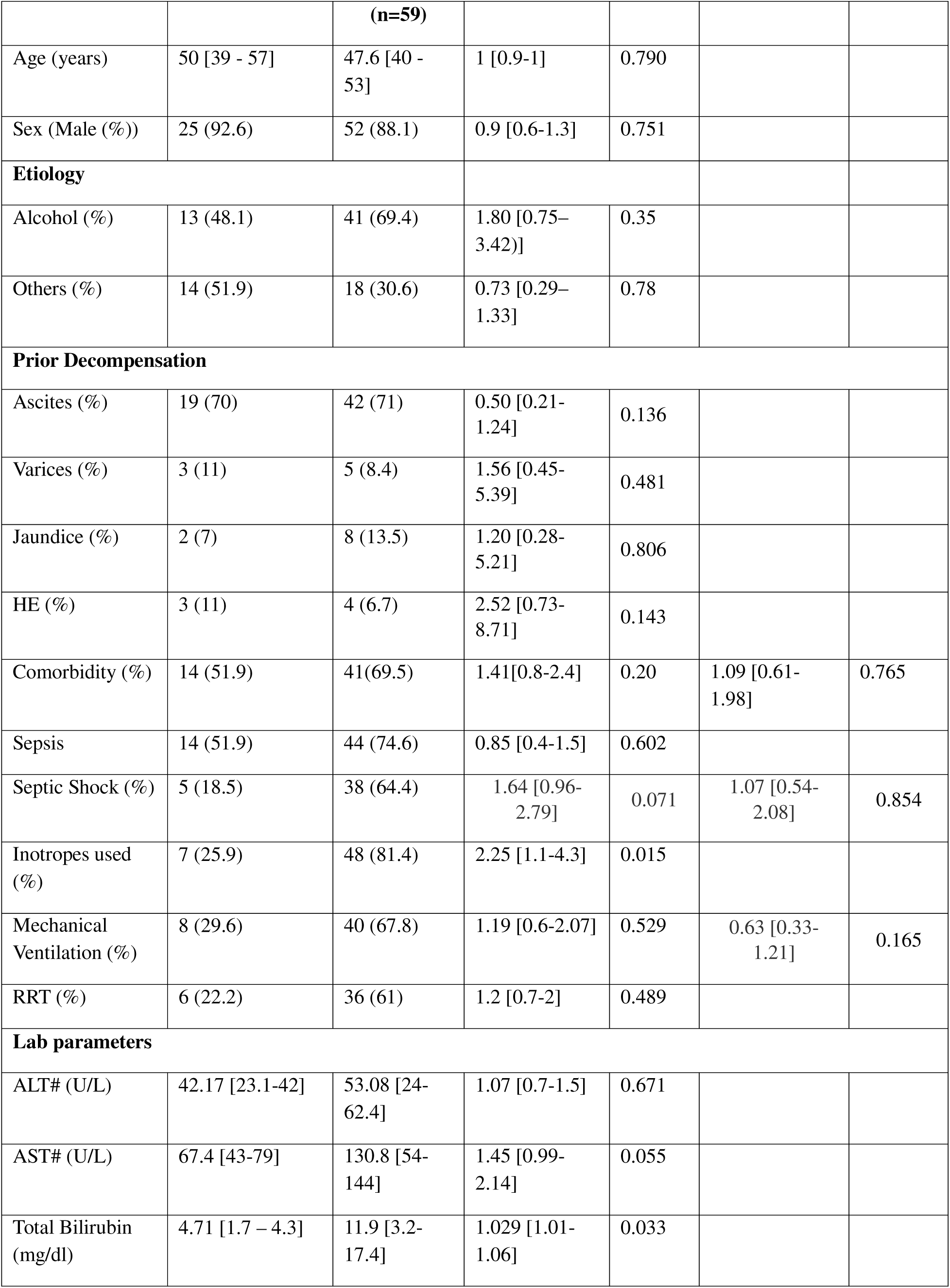
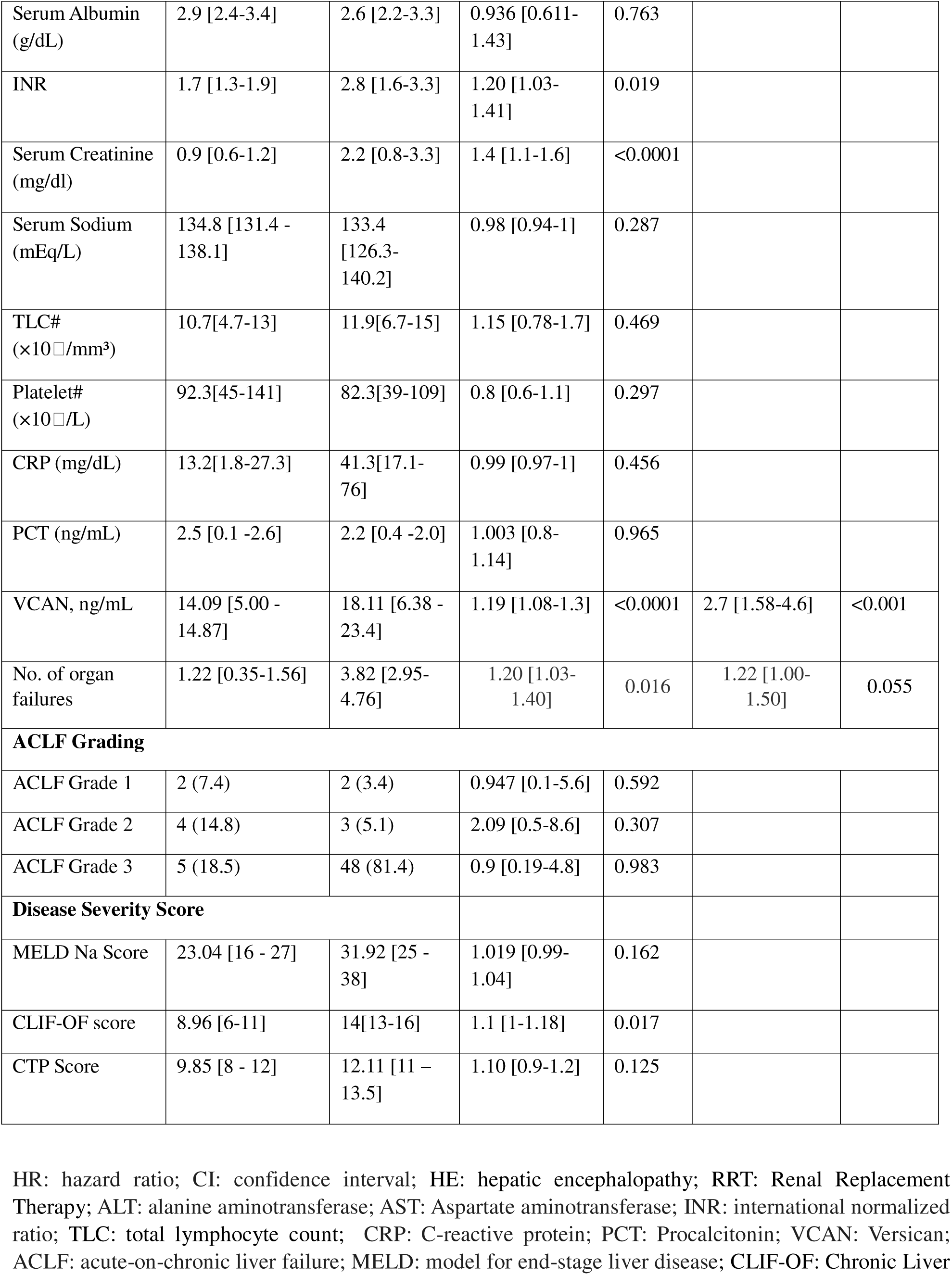
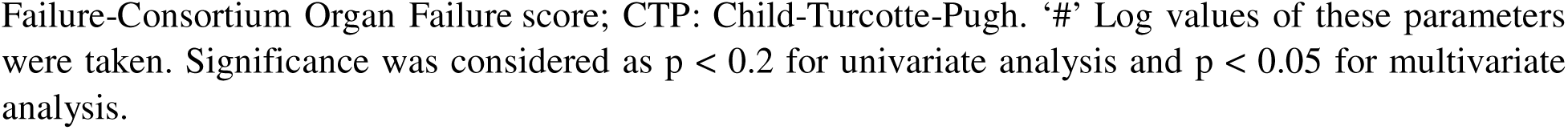
Univariate and multivariate Cox regression analysis of 28-day mortality in critically ill cirrhotic patients.

During the 28-day follow-up, 42 patients (48.8%) died in the cirrhosis group, mainly from septic shock or infectious complications. 15 patients (17.4%) underwent liver transplantation, and 2 (2.3%) received TIPS. Non-survivors showed significantly higher AST, bilirubin, creatinine, INR, leukocyte count, CRP, PCT, MELD-Na, and infection rates, compared with survivors. Importantly, VCAN levels were markedly elevated in non-survivors (18.11 ng/mL) compared with survivors (14.09 ng/mL, p < 0.0001; **Fig. 6C**). Among cirrhotic patients, VCAN was also higher in those with sepsis (17.2 ng/mL) than without (15.9 ng/mL, p=0.0252; **Fig. 6D**), and ascitic fluid VCAN levels were increased in patients with SBP (p=0.035 **Fig. 6E**). On univariate Cox regression analysis, VCAN levels were significantly associated 28-day mortality, alongside these parameters: comorbidity, septic shock, organ failure, and severity scores (each p < 0.02). In the multivariate model, VCAN level (HR: 2.7, 95% CI: 1.58 - 4.6, p < 0.001) came out as an independent predictor of 28-day mortality along with organ failure and mechanical ventilation. The ROC curve analysis further confirmed the discriminative power of plasma VCAN for predicting 28-day mortality (AUC = 0.841, 95% CI: 0.744 - 0.937, p < 0.0001; **Fig. 6F**). A cutoff of 15.9 ng/mL of plasma VCAN levels provided 83% sensitivity and 71% specificity. Based on this threshold, patients were stratified into high and low VCAN groups. Kaplan–Meier survival analysis revealed significantly reduced survival in the high-VCAN group (χ² = 9.494, df = 1, p = 0.002; **Fig. 6G**), with 87.5% mortality in patients with VCAN >15.9 ng/mL compared with 33% in those below the cutoff. Because liver transplantation and TIPS procedures represent alternative outcomes that preclude mortality, we also incorporated them in a competing-risk regression model to validate the robustness of the association. In this model, VCAN remained a significant predictor of adverse outcome, with a SHR of 1.22 (95% CI: 1.12–1.33, p < 0.001, **Table 2**; **Fig. 6H**), indicating that higher VCAN levels significantly increased the likelihood of mortality. Collectively, these findings demonstrate that elevated plasma VCAN levels are independently and strongly associated with increased short-term mortality in critically ill patients with cirrhosis.

**Table 2:**
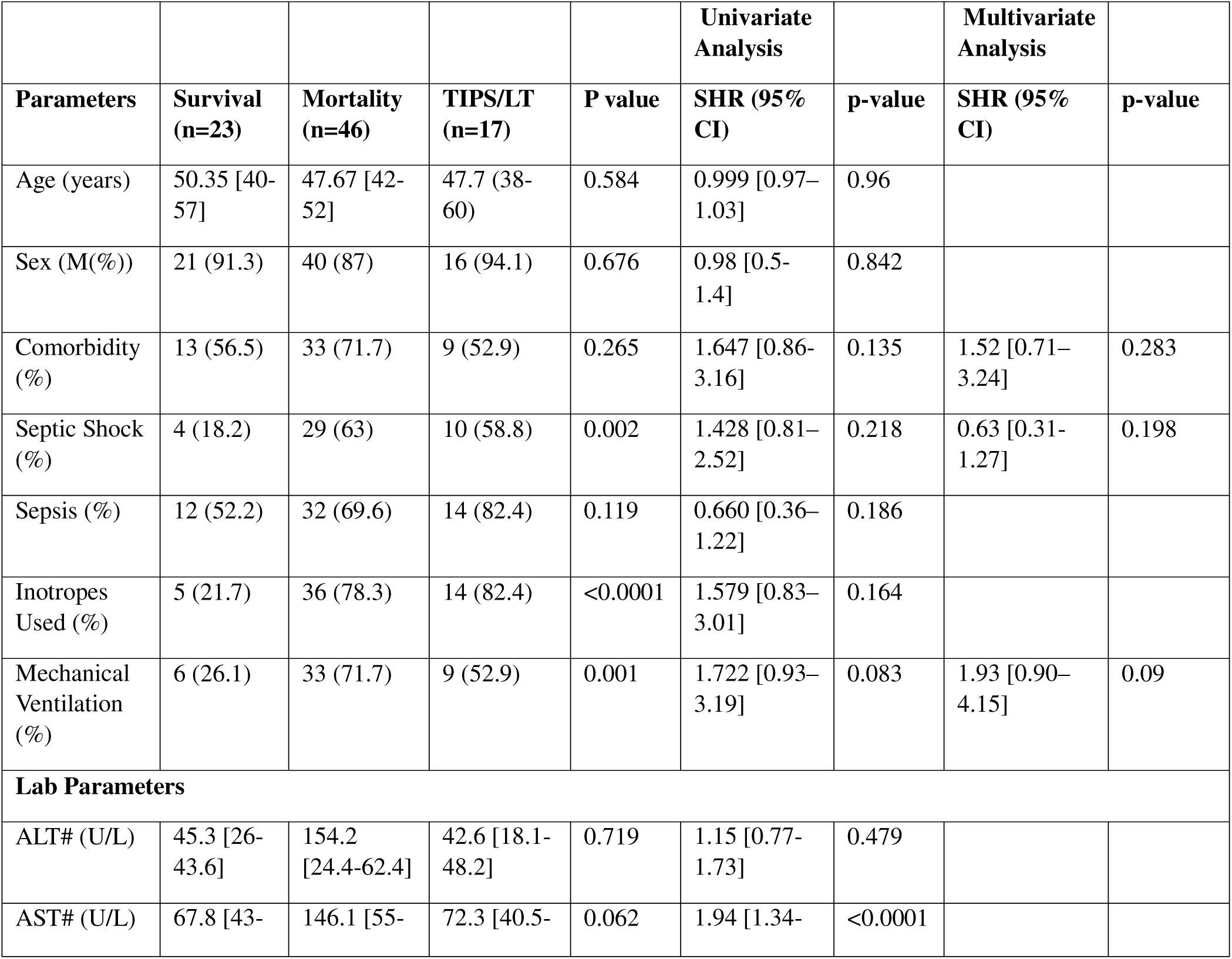
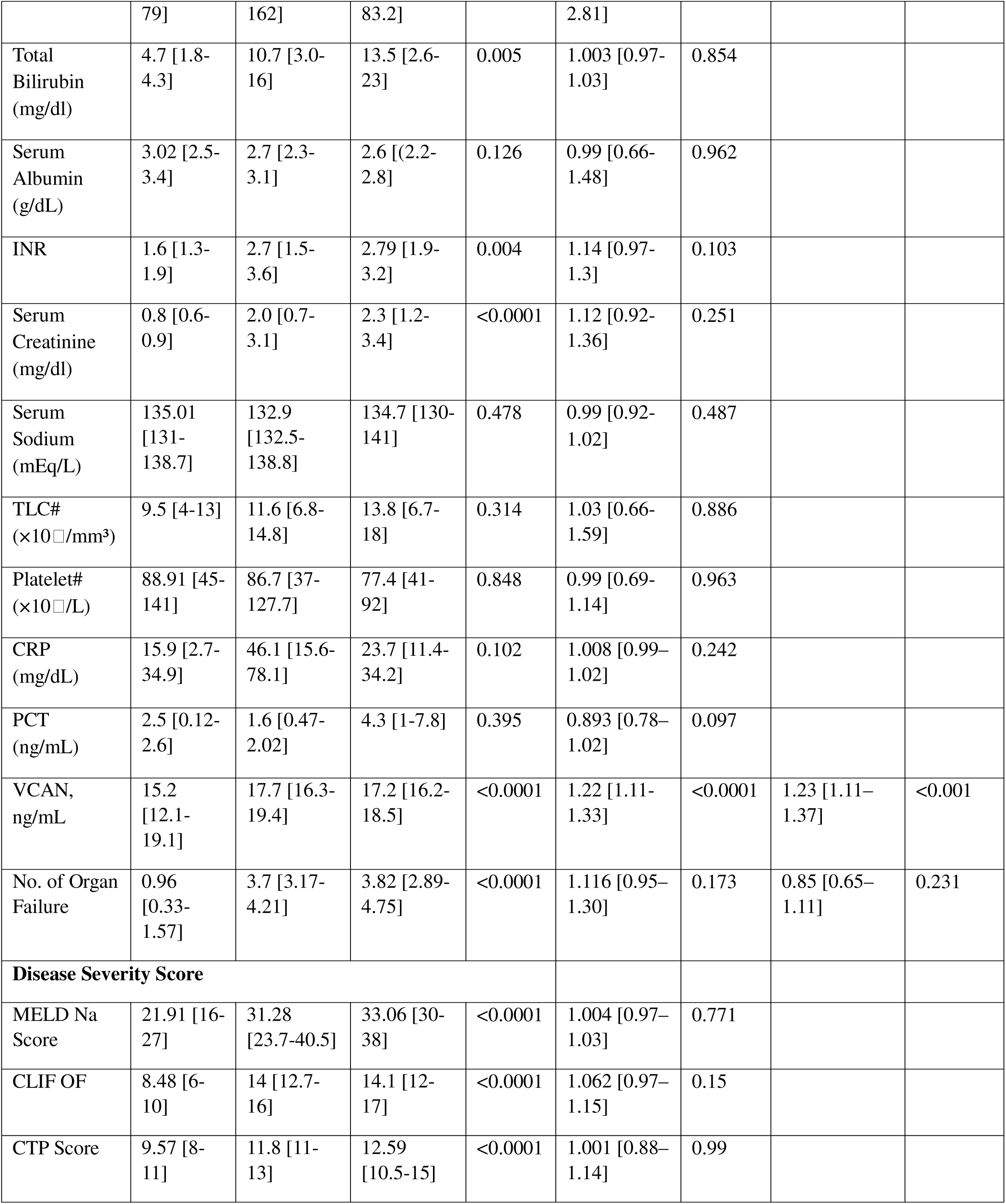
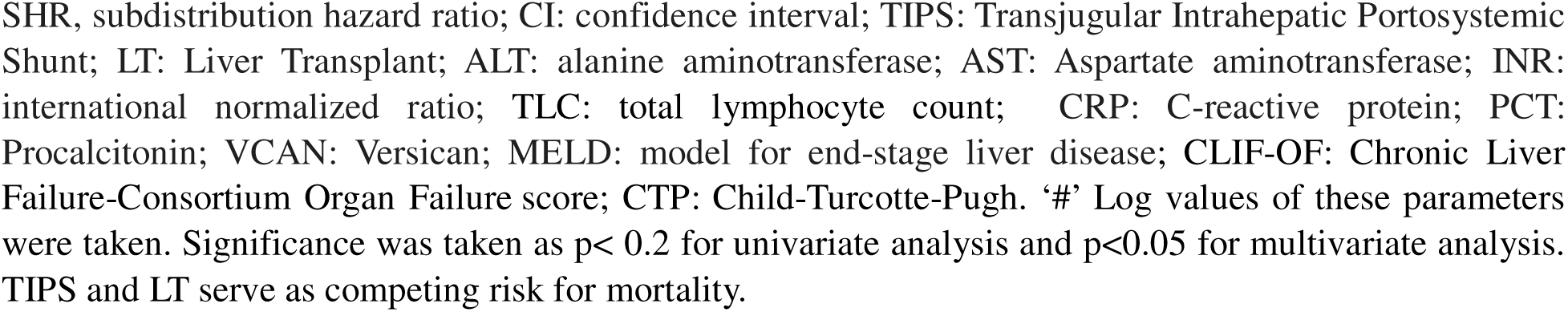
Results of Univariate and Multivariate Fine and Gray Model for Risk Factors of 28-Day mortality in cirrhosis patients.

## Discussion

Our findings reveal that in cirrhosis, MLN are no longer effective microbial gatekeepers due to both fibrotic remodeling and immune cell dysfunction. We identify up-regulation of the ECM protein, VCAN, within the cirrhotic MLN stroma as key driver of this impairment. Elevated VCAN expression in MLN disrupts T cell trafficking, suppresses CD4 T cell activation, enhances immunosuppression, and facilitates systemic bacterial dissemination, collectively leading to adverse clinical outcomes. These results not only uncover a novel mechanism of MLN dysfunction in cirrhosis but also underscore VCAN as a potential biomarker in infection-related mortality in cirrhosis patients.

In concordance with previous studies, we demonstrate that healthy animals efficiently contained gut bacteria within MLN, preventing systemic spread [13,14]. In contrast, cirrhotic rats showed marked bacterial accumulation in MLN, portal blood, and peripheral organs. Surgical removal of MLN in control rats caused bacterial dissemination to the extraintestinal organs, including the liver, spleen, and lungs, confirming MLN as a crucial barrier. Consistently, MLN removal in cirrhotic animals exacerbated BT, endotoxemia in distant organs, i.e., spleen and lungs, with increased liver fibrosis, and reduced survival, highlighting immune barrier failure as a direct contributor to disease progression. In a previous study, authors reported that *Salmonella* infection post MLN removal increased to 100-fold in systemic circulation post 2 days in otherwise resistant mice, with microbial presence in the liver and spleen as well. This was accompanied by elevated liver injury markers, TNF-α, and IFN-γ cytokines [15]. Together, these findings underscore that MLN integrity is vital to limit BT, consistent with clinical observation linking gut-derived infections to morbidity and mortality in cirrhosis [1,16].

Under physiological conditions, MLN serve as central hubs for antigen presentation and T cell priming, ensuring a delicate equilibrium between immune tolerance to commensals and defense against pathogens [17,18]. This equilibrium is perturbed in cirrhosis, where MLN undergoes profound structural expansion and remodeling in response to chronic liver injury [19–21]. In our previous studies, we have reported aberrant lymphangiogenesis and lymphatic dysfunction in the mesentery and MLN, which facilitate BT, fueling systemic infections and inflammation [22]. Here, we performed a detailed analysis of immune cell subsets in MLN in advanced liver disease. Our results revealed compartmentalized immune alterations within MLN and systemic circulation in cirrhotic animals post-LPS challenge. Elevated frequencies of activated CD4 T cells, antigen-presenting DCs, and macrophages within cirrhotic MLN suggest a hyperactive local immune response, likely triggered by ongoing bacterial stimuli. However, this was also accompanied by increased CD8^+^CD25^+^ Treg cells and TGF-β in MLN and systemic circulation, reflecting a microenvironment skewed toward immune suppression. These divergent immune responses suggest that local MLN activation alongside systemic immune paralysis in cirrhosis may impair pathogen clearance and increase bacterial infection. A similar finding revealed that CXCR6 CD69 CD8 T cells within the ascitic fluid of cirrhotic patients exhibit a hyperactivated yet functionally impaired phenotype, distinct from circulating and hepatic counterparts, indicating compartment-specific immune dysregulation [23]. Previous study demonstrated that bacterial accumulation in the MLN drives Th1 differentiation and monocyte-driven TNF-α production in cirrhotic rats, resulting in re-circulation of these activated immune cells into the blood, promoting systemic inflammation [24].

In our study, the role of MLN in shaping systemic immunity was further supported by the observation that MLN resection in healthy rats reduced systemic activated CD4 T cell frequency. In cirrhotic rats, however, we observed less pronounced effects after MLN removal, likely because baseline MLN dysfunction in cirrhosis already impairs immune regulation. This highlights the substantial contribution of MLN in sustaining systemic T cell activation under physiological conditions, but their compromised function in cirrhosis attenuates this immunostimulatory capacity. Also, MLN resection led to minimal compensatory changes detected in the spleen, especially in cirrhosis, underscoring the non-redundant role of MLN.

Proteomic profiling revealed key mechanistic aspects of MLN dysfunction in cirrhosis. Notably, antimicrobial defense proteins such as β-defensins, nuclear receptor coactivator 3, and doublecortin were markedly downregulated, indicating impaired innate immune responses. In contrast, proteins associated with ECM remodeling and immune activation, including VCAN, SLIT-ROBO Rho GTPase-activating protein 2, CCR10, and granzyme, were strongly upregulated, suggesting heightened immune activation and tissue remodeling. VCAN, a large chondroitin sulfate proteoglycan abundant in the ECM of soft tissues, plays a dynamic role in immune regulation due to its highly charged glycosaminoglycan side chains. By interacting with key innate immune mediators, such as hyaluronan, TLR-2, cytokines, chemokines, and growth factors, VCAN can modulate inflammatory responses, either amplifying or restraining them depending on the microenvironmental context [25–27].

The liver is known to handle bacterial load originating from the portal and mesenteric circulation [28]. In our BDL models, bacterial load was detectable in MLN at week 3 post BDL, indicating that systemic spread of bacteria was handled by the liver firewall during this stage. However, post 4 weeks, bacterial load was detected in large numbers in MLN, liver, and other organs as well, clearly depicting the shutdown of both liver and MLN defense systems, and further in other organs. Also, we observed a progressive increase in VCAN that correlated with fibrosis severity in both liver and MLN, bacterial load, and elevated TGF-β in MLN, linking VCAN expression with cirrhosis progression and BT burden. Also, VCAN levels were significantly elevated in MLN with detectable bacterial load compared to those without.

Functional analyses further highlighted VCAN’s immunomodulatory role under both *in vitro* and *in vivo* conditions. We first identified cirrshotic FRCs as the primary source of VCAN in MLN. rVCAN inhibited T cell activation by increasing CD25^+^ Treg cell populations. It also impaired T cell migration across FRC layers and enhanced Treg-mediated suppression of activated T cells. Blocking CD44 on T cells restored this migration, consistent with prior evidence that VCAN-CD44 interactions regulate T cell trafficking [12]. A previous study reported that FRCs make intimate contact with T cells via CD44, mediate the induction of CD8 T cell tolerance, and possess potent suppressive capability [29]. Thus, excessive VCAN deposition likely creates a dense matrix that restricts T cell movement and fosters immunosuppressive conditions within cirrhotic MLN, facilitating bacterial persistence. *In vivo*, siRNA-mediated VCAN knockdown in cirrhotic rats restored T cell migration to systemic circulation with effective T cell activation and reduced TGF-β levels in both MLN and systemic circulation. VCAN knockdown decreased bacterial load and, importantly, fibrosis within the MLN. However, we did not observe a reduction in bacterial burden in other organs after VCAN knockdown, as it depends on broader immune mechanisms. Previous studies have demonstrated that transient knockdown of VCAN in hepatic stellate cells reduced fibrogenesis markers and cell proliferation, suggesting its role in modulating tissue remodeling and inflammation [11]. Our study for the first time illustrates a mechanistic link between VCAN-mediated fibrotic remodeling in MLN, impaired adaptive immunity, and susceptibility to bacterial infections, underscoring its potential as a therapeutic target in cirrhosis.

In cirrhosis, progressive liver fibrosis drives ECM remodeling, resulting in the release of ECM fragments into circulation. Circulating ECM proteins such as hyaluronic acid, procollagen type III N-terminal peptide, and tissue inhibitor of metalloproteinase-1 have established roles in noninvasive fibrosis assessment and prognosis [30–32]. Hence, we next investigated plasma VCAN levels in clinical settings in patients with decompensated cirrhosis. Elevated plasma VCAN levels were clearly seen in cirrhotic patients with bacterial infections. Its increase in patients with bacterial infection may reflect intensified inflammatory activation and impaired microbial clearance within lymphoid tissue. In our multivariate analysis, VCAN, along with organ failure and mechanical ventilation, remained significant predictors of short-term mortality. The ROC analysis confirmed the discriminative power of VCAN for 28-day mortality, with an AUC of 0.841 (95% CI: 0.744-0.937), which is comparable with the established markers such as vWF and CLIF-C-ACLF score in a previous study [33].

A plasma VCAN cutoff of 15.9 ng/mL provided high sensitivity (83%) and specificity (71%), to stratify patients for mortality risk. Survival analysis strengthened this prognostic significance, with markedly reduced survival in patients with high VCAN levels (87.5% vs. 33%), indicating that VCAN can effectively differentiate patients at high risk of early death. Importantly, these findings remained consistent even after accounting for competing outcomes such as liver transplantation and TIPS, where VCAN retained its predictive value in the competing-risk regression model (SHR = 1.22, 95% CI: 1.12–1.33, p < 0.001). In this model, mechanical ventilation also emerged to be a borderline significant factor. This consistency strengthens the reliability of VCAN as a biomarker of adverse prognosis, independent of treatment-related confounding. Our results align with emerging evidence that ECM remodeling markers are critical in disease progression. ECM protein, PRO-C3, and PRO-C6, which reflect interstitial collagen formation and fibroblast activity, were reported to be elevated in ACLF and linked to fibrosis and inflammation [32]. The selective increase of interstitial ECM formation markers in ACLF supports the concept that fibroblast-driven ECM remodeling plays a key role in organ functional decline and containment of bacterial infections. CRP level and MELD score are unreliable in sepsis with cirrhosis due to impaired hepatic synthesis [34,35]. In one of the studies, authors have reported that sTREM-1, a myeloid-derived protein, better predicts infection and 90-day mortality in cirrhotic patients (AUROC=0.733, CI:0.59-0.86, p=0.0027) and (HR:2.94, CI:1.009-8.5, p=0.0048) [36]. We suggest that extrahepatic proteins like VCAN in MLN may serve as more reliable biomarkers of disease progression in advanced cirrhosis. This major limitation of this study is a relatively small sample size and a single-center design. In addition, VCAN levels were measured at baseline; tracking their changes over time could offer deeper insights into their role in disease progression.

In summary, our data emphasize that MLN are vital gatekeepers in preventing bacterial dissemination in liver cirrhosis, with their dysfunction significantly contributing to systemic endotoxemia and disease progression. By establishing a mechanistic link between stromal matrix remodeling, particularly VCAN upregulation, and impaired T cell activation and migration, our findings reveal how excessive ECM deposition fosters an immunosuppressive niche that permits bacterial persistence. The strong association between circulating VCAN levels and infection-related mortality underscores its potential as a biomarker. Future studies should explore VCAN-targeted interventions and validate their impact on immune restoration and clinical outcomes in cirrhosis.

## Declaration of generative AI and AI-assisted technologies in the manuscript preparation process

During the preparation of this work the authors used Perplexity AI in order to improve clarity. After using this tool/service, the authors reviewed and edited the content as needed and takes full responsibility for the content of the published article.

## Statements and Declarations

### Data Availability

All the data supporting the findings of this study are available within the article and its supplementary information files and from the corresponding authors upon reasonable request.

### Author Contributions

SK and PJ conceptualized and designed the work. PJ, AS, DJ, BS, and NC collected the data. PJ, AS, BS, VK, RM, GK, and SK analyzed and interpreted the data. PJ and SK drafted the article. PJ, BS, SK, AM, DMT and SKS did critical revision of the article. All authors read and approved the final manuscript.

---

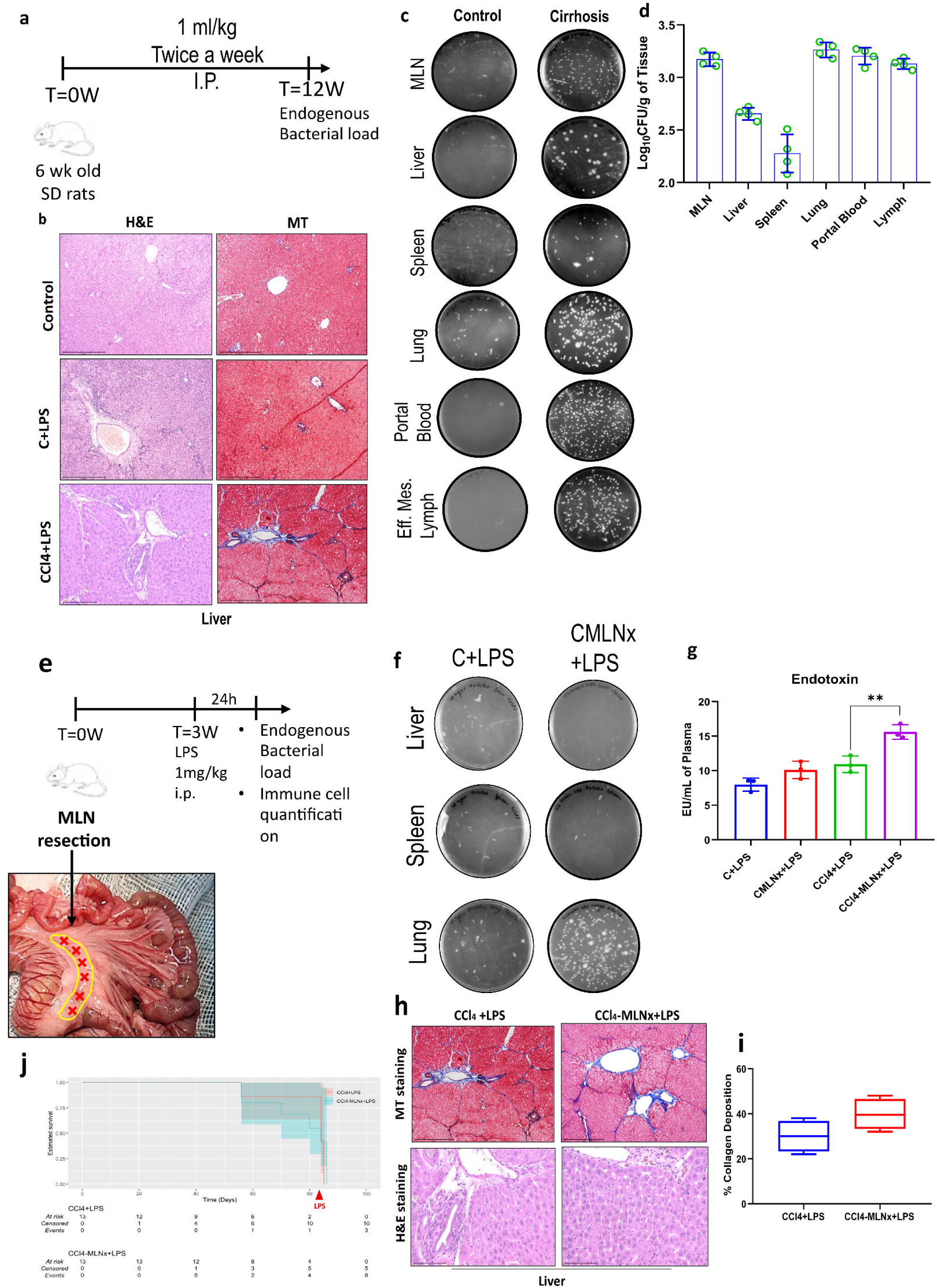

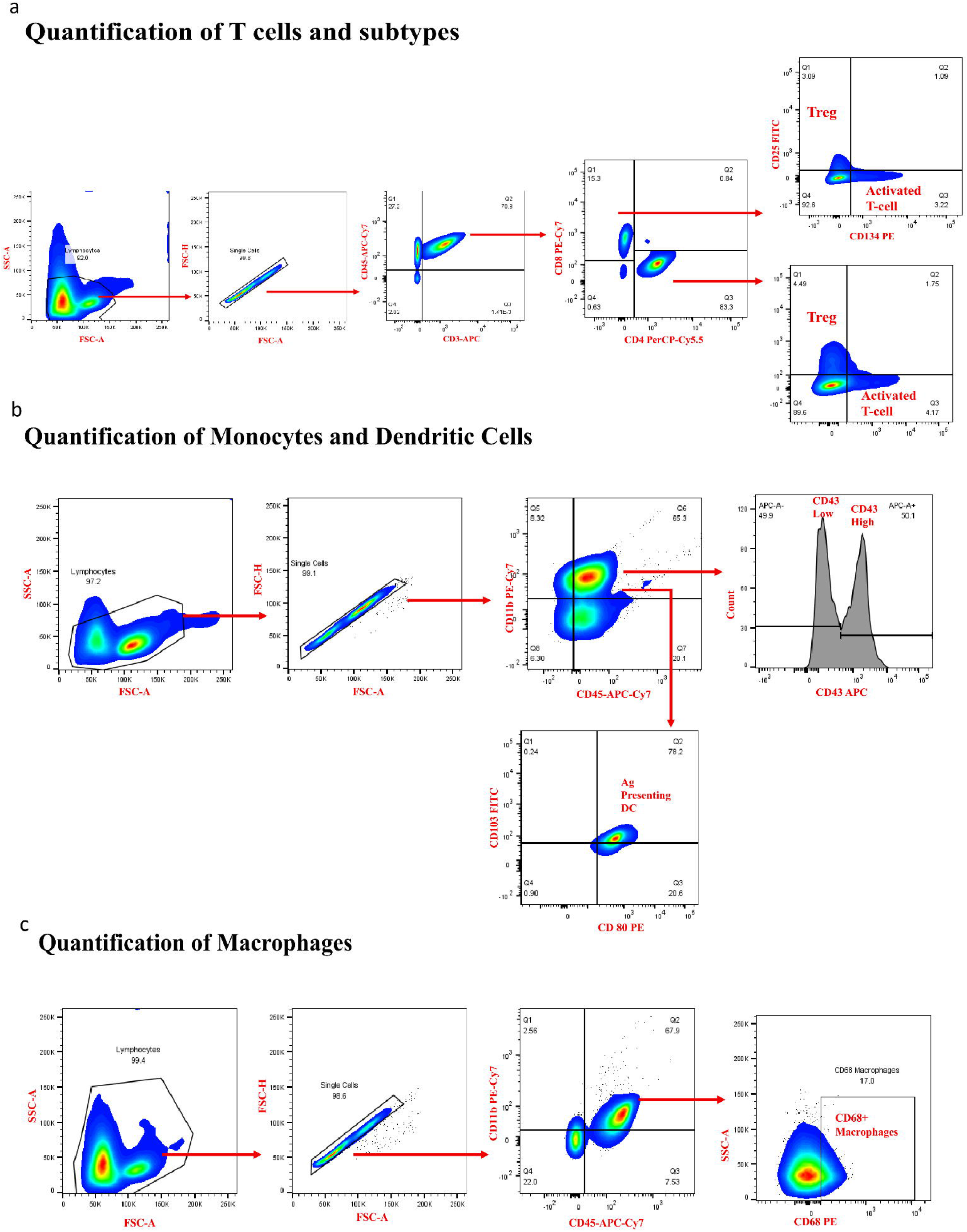

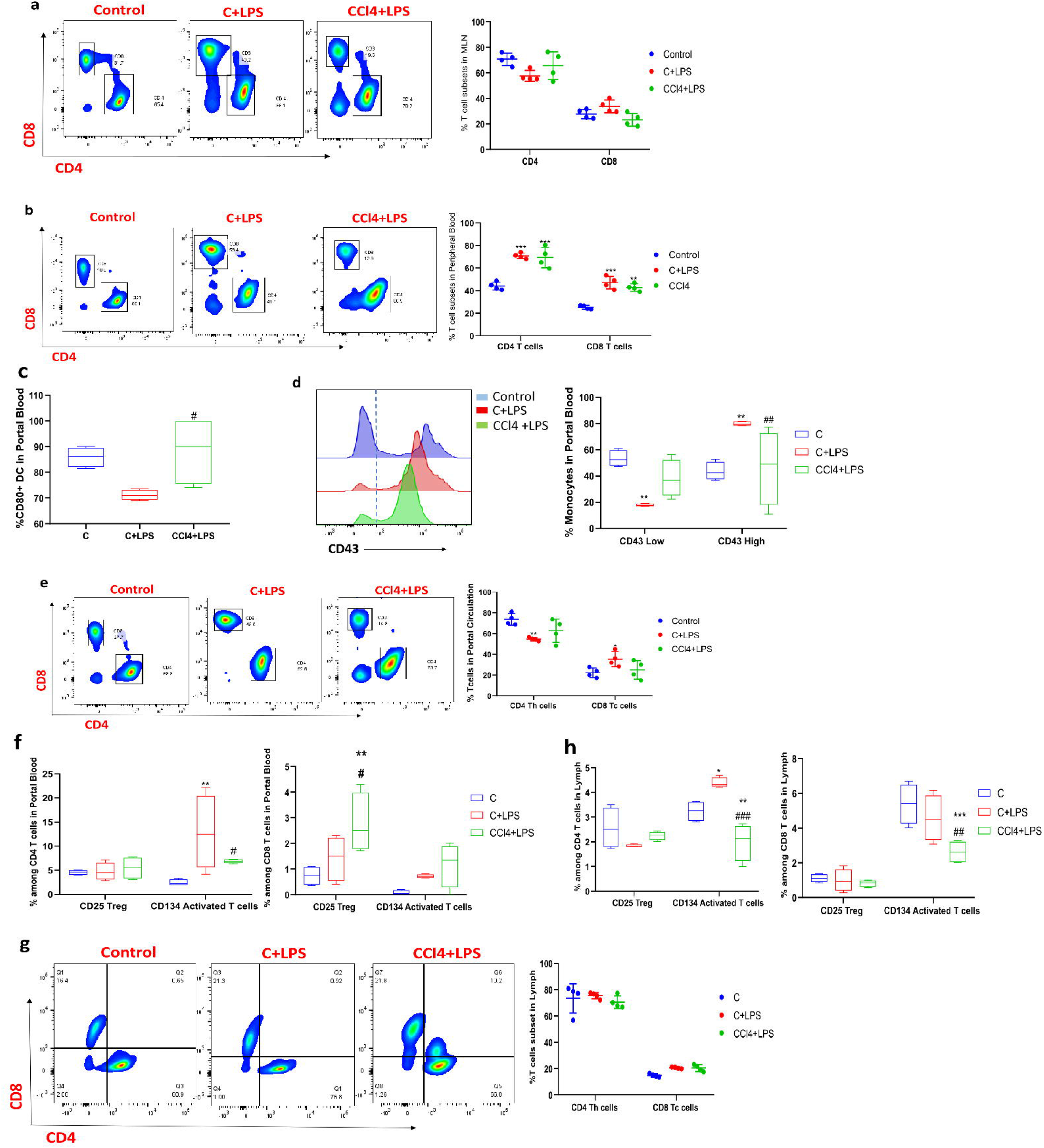

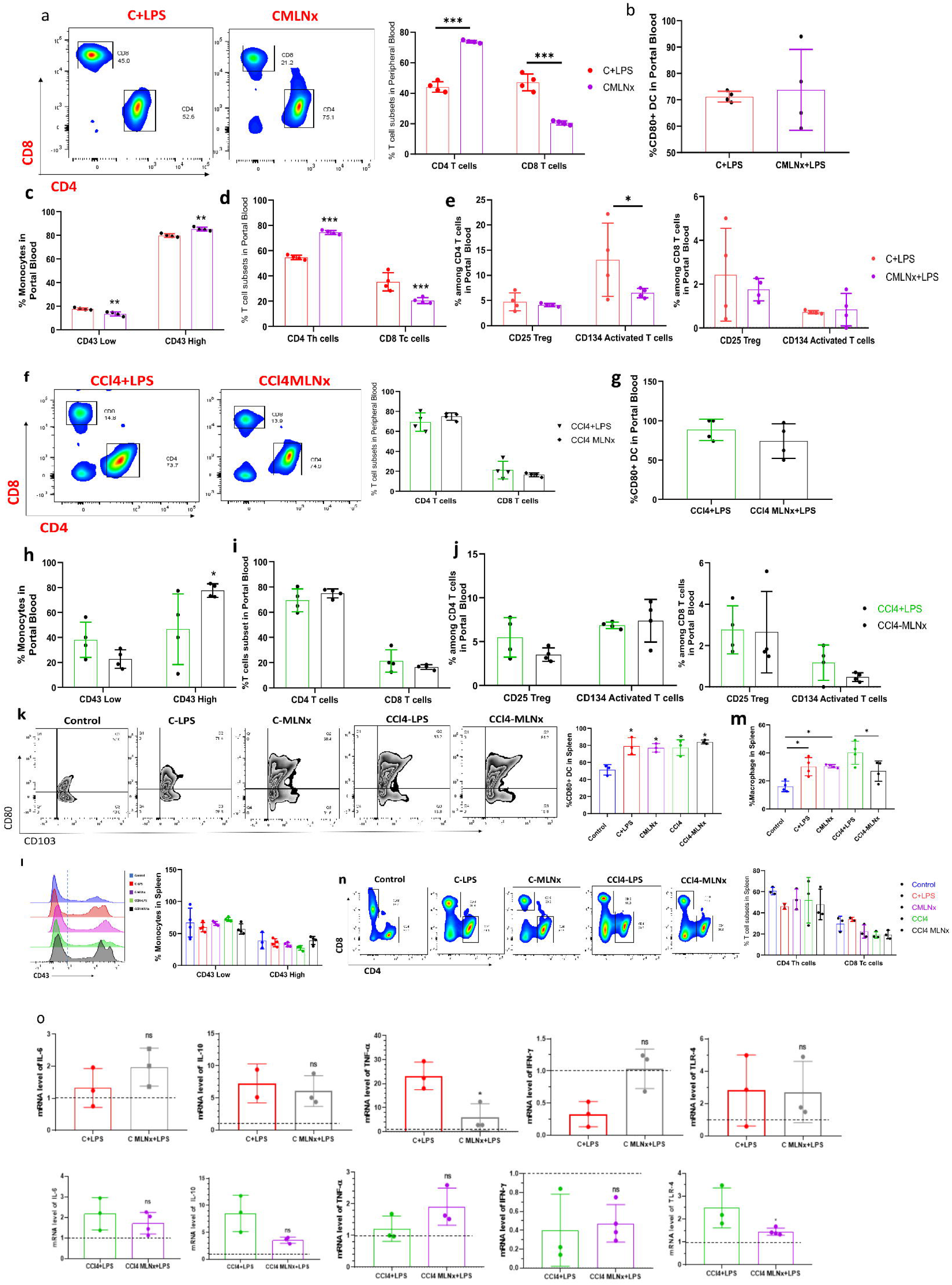

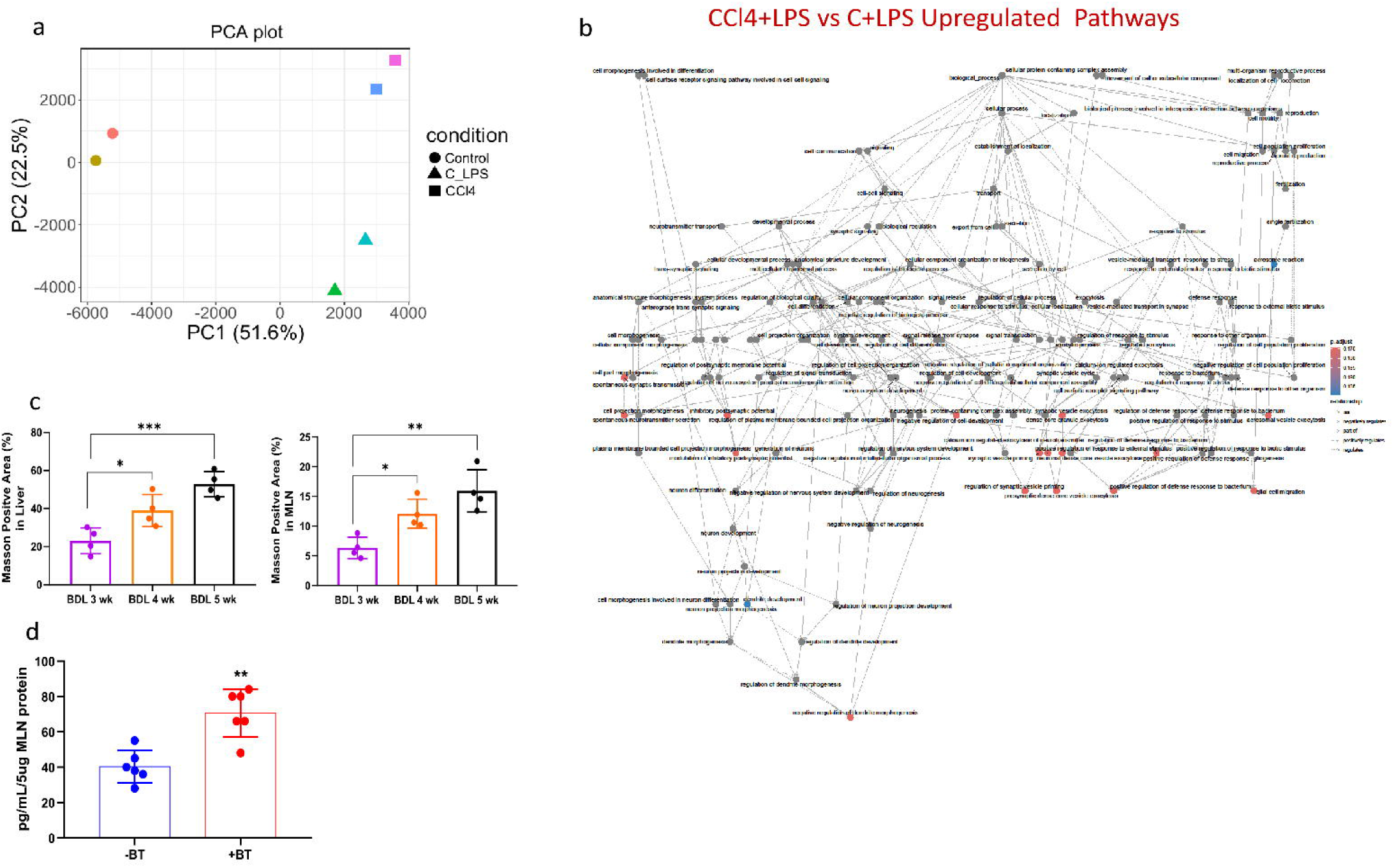

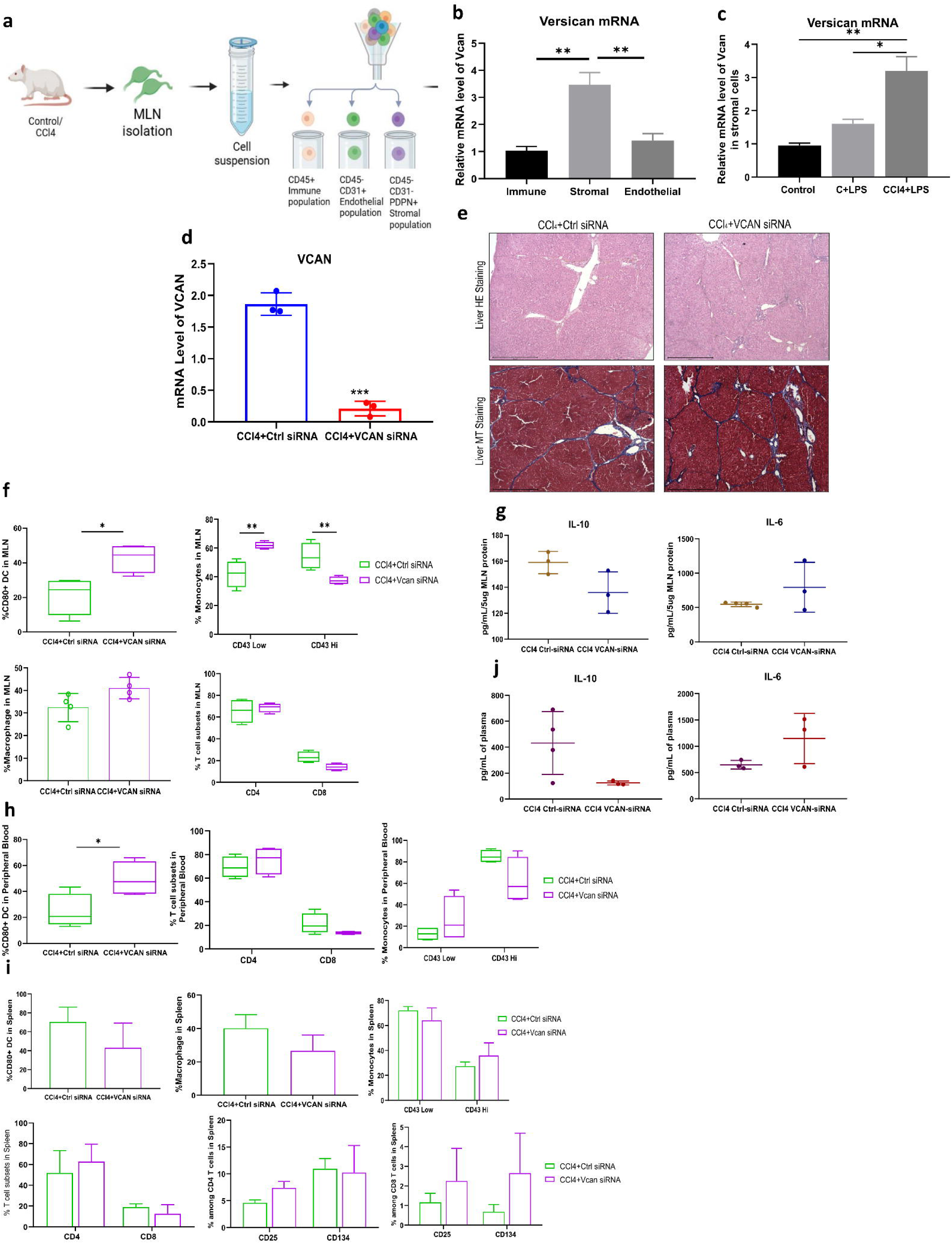

