## Supplementary material for "Increased versican and fibrosis in mesenteric lymph nodes disrupts immune surveillance and drives systemic bacterial dissemination in cirrhosis": Supp file

**Table of Contents:**

Material and methods………………………………………………………………………………..2

Supplementary figure legends……………………………………………....…….............................9

Supplementary Table…………………………………………………………………………….….11

References………………………………………………………………………………………… 19

**Methods**

**Animal Experiments and Model Preparation**

All animals were provided humane care in accordance with the *Guide for the Care and Use of Laboratory Animals* as outlined by the National Academy of Sciences and published by the National Institutes of Health (NIH publication 86-23, revised 1985). The study was approved by the Animal Ethics Committee of ILBS, New Delhi, in accordance with standard ethical guidelines (Ethics Protocol No: IAEC/ILBS/21/15). Rats were housed in a controlled environment at 22±3°C with a 12-hour light/dark cycle, and were given food and water ad libitum. The study used 6-week-old male Sprague-Dawley (SD) rats weighing 200–250 g. Liver fibrosis was induced via intraperitoneal (i.p.) injection of a 1:1 mixture of CCl_4_:olive oil at a dosage of 1.0 mL/kg body weight, administered twice a week for 12 weeks. Phenobarbitone was added to the drinking water. Rats were assigned to three experimental groups: control (no treatment or injection), C+LPS (1 mg/kg LPS, i.p. injection), and CCl_4_+LPS (12 weeks of CCl_4_ administration followed by a single 1 mg/kg LPS injection). 24 hrs post-LPS injection, the rats were sacrificed, and tissues were collected for subsequent analysis. A bile duct ligation (BDL) animal model in rats was prepared by ligating the bile duct as previously described (Juneja et al. 2023). Animals were sacrificed at 3, 4, and 5 weeks post-surgery, and tissues were collected for subsequent analysis.

**Resection of MLN**

Under isoflurane anesthesia, the procedure was conducted under sterile conditions. The body temperature was carefully maintained at 37°C throughout the surgery. A laparotomy was performed, and the intestine was gently exteriorized on moist gauze. Saline was continuously applied to the intestine to prevent tissue desiccation. The mesenteric lymph nodes (MLN) were exposed, and the remaining intestine was covered with sterile gauze to maintain a sterile environment. Using fine, pointed forceps, the tissue surrounding the MLN from the duodenum to the ileum was carefully dissected. The MLN were then extracted with the aid of blunt, rounded forceps. Once all the MLNs were removed, the intestine was repositioned, and the abdominal incision was closed with sutures. Post-operative care was administered for three days, and the animals were allowed to rest for three weeks to facilitate revascularization at the surgical site. Subsequently, the animals were randomly divided into two groups: CMLNx (LPS injected i.p. at 1 mL/kg) and CCl_4_-MLNx (CCl_4_ administered for an additional 12 weeks, followed by a single i.p. injection of LPS at 1 mg/mL). Twenty-four hours after LPS administration, the animals were sacrificed, and tissue samples were collected for further analysis.

***In vivo* *VCAN* siRNA delivery in rats:**

**siRNA**-*In vivo* ready synthetic siRNA was purchased from GeneX India Bioscience, and the most efficient sequence at knocking down specific targets was used *in vivo*. Scrambled control siRNA (sequence: sense 5' GCATGAAAGTCCCTAATAA 3'); *VCAN* specific siRNA (sequence: sense 5′-CUCAAGCGAUAUACAAUGA -3′).

**siRNA/*In vivo* JET-PEI complex preparation**: *In vivo* JET-PEI (cat. no 201-10G, Polyplus-transfection S.A, Illkirch, France) was used according to the manufacturer's protocol. Preparation of the siRNA/*in vivo* JET-PEI complex in N/P ratio 8 was performed as follows. The siRNA was first diluted in 100 µL of RNase-free water to reconstitute at a 1 mg/mL concentration. It was then mixed with 100 µl of 10% glucose to prepare solution A. 16 µl of *In vivo* JET-PEI in 84 µl water was mixed with 100 µl of 10% glucose to prepare solution B. Solution A, and B were mixed, vortexed, and incubated for 15 minutes at RT. For the *in vivo* experiments, 80 µg of siRNA was injected per rat in three doses on alternate days.

**Assessment of Bacterial Load**

All experiments were conducted under sterile conditions. Rats were anesthetized with an intraperitoneal (i.p.) injection of ketamine hydrochloride (60 mg/kg) and midazolam (3 mg/kg), followed by shaving and disinfection of the skin with alcohol. After performing a midline laparotomy, MLN were dissected, excised, and weighed in a sterile environment. Tissue samples from the liver, spleen, and lungs were also collected and weighed. All tissue specimens were homogenized in phosphate-buffered saline (100 mg per 100 µL), and the resulting suspension was cultured on luria broth (LB) agar. The culture was incubated for 24 hrs, and bacterial colonies were imaged and counted using ImageJ software. Colony-forming units (CFU) were calculated and plotted on Log 10 scale. For bacterial translocation (BT) studies, exogenous GFP-labelled *Salmonella typhi* (10^9^ bacterial cells) were administered by gavage, and rats were sacrificed 48 hrs post-gavage. Organs were harvested in the same manner and plated on LB agar supplemented with ampicillin for selection of GFP-labeled bacteria, allowing for the growth of GFP-translocated bacterial strains.

**Sample Preparation for Proteomics**

Tissue samples for proteomic analysis via mass spectrometry are promptly harvested, flash-frozen in liquid nitrogen, and stored at -80°C until further processing. The frozen tissue is homogenized in a pre-chilled lysis buffer containing 8 M urea, 1 M thiourea, and protease/phosphatase inhibitors. The homogenate is then centrifuged at 12,000 x g for 20 minutes at 4°C to remove cellular debris, and the protein-containing supernatant is collected. Protein concentration is determined using BCA. To ensure complete protein denaturation, proteins are reduced with DTT or TCEP and incubated at 37°C, then alkylated with iodoacetamide to prevent cysteine oxidation. Proteins are digested overnight with trypsin at a 1:50 enzyme-to-substrate ratio. The resultant peptides are desalted using C18 columns to remove contaminants, then concentrated by lyophilization. Peptide concentration is measured, and the peptides are reconstituted in 0.1% formic acid for analysis. The peptides are subjected to LC/MS for detailed proteomic profiling.

**Histology**

Tissue sections (7-µm thick) fixed in 10% formalin are first deparaffinized by immersion in xylene, followed by rehydration through a descending ethanol series. For H&E staining, sections are stained with hematoxylin, differentiated in acid alcohol, and blued in running tap water. The sections are then stained with eosin, dehydrated through an ascending ethanol series, cleared in xylene, and mounted with a coverslip. For Masson's Trichrome (MT) staining, after deparaffinization and rehydration, sections are stained with hematoxylin, followed by acid fuchsin, and differentiated in phosphomolybdic acid. Sections are then stained with aniline blue, differentiated in 1% acetic acid, dehydrated, cleared in xylene, and mounted with a coverslip. MT staining in tissue samples was quantified using ImageJ and plotted using area% in graphs.

**Isolation of cells from rat MLN nd blood for *in vitro* culture and flow cytometry**

For cell isolation from MLN, rats were anesthetized with an i.p. injection of ketamine hydrochloride (60 mg/kg) and midazolam (3 mg/kg). The MLN were carefully excised and placed in DMEM supplemented with 1% antibiotic and antimycotic solutions. Using sterile scissors, the tissue was finely minced into approximately 1 mm³ pieces. To facilitate *in vitro* digestion, 0.25% collagenase IV, prewarmed to 37°C, was prepared. The minced tissue was resuspended in the enzyme solution and incubated with constant shaking to achieve optimal digestion. After 10-15 minutes of digestion at 37°C, the tube was transferred to a biosafety cabinet, and the digested tissue suspension was filtered through a 70 μm strainer into a sterile 50 ml tube. An equal volume of sterile DMEM was added to the strainer to neutralize the enzyme activity. The resulting cell suspension was then centrifuged at 1200 rpm for 5 minutes at 4°C and washed twice with PBS. If necessary, RBC lysis was performed. Following cell counting, the cells were stained with CD31-PE (BD Biosciences, 555027), PDPN-APC (BioLegend, 250906), and CD45-APC-Cy7 (BioLegend, 202216) antibodies. To isolate specific populations, FRCs, immune cells, and endothelial cells were sorted using FACS. The sorted cells were collected in the media. After a final cell count, the cells were plated in a culture dish containing complete DMEM supplemented with 1% antibiotic and 10% FBS. For flow cytometry, 1 million cells were incubated with antibodies specific for T cell subsets, dendritic cells (DC), monocytes, and macrophages for 30 minutes in the dark at RT. A total of 2-4 lakh events were acquired for each experiment.

For isolation of immune cells from blood, 1X RBC lysis buffer was used. For 1 ml of blood, 9 ml of buffer was added. The cell pellet free from RBC was washed with PBS to remove excess RBC lysis buffer. Cells were then counted using a hemocytometer, and 1 million cells were incubated with antibodies for immune cell quantification. A total of 2-4 lakh events were acquired for each experiment

**RNA extraction and RT-PCR**

RNA extraction and RT-PCR were performed on excised mesenteric tissues stored in an RNA buffer. Total RNA was isolated using a Nucleopore kit, according to the manufacturer's instructions. RNA was quantified at 260/280 nm using a Nanodrop 2000 spectrophotometer (Thermo Scientific). First-strand cDNA was synthesized from 1µg of total RNA using reverse transcriptase (Thermo Fisher Scientific Verso cDNA synthesis kit) according to the manufacturer's instructions. Quantitative real-time PCR was performed using SYBR Green PCR Master Mix (Fermentas Life Sciences) on a ViiA7 Real-Time PCR System (Applied Biosystems, USA). The following cycling parameters were used: start at 95 °C for 5 min, denaturation at 95 °C for 30 s, annealing at 60 °C for 30 s, elongation at 72 °C for 30 s, and a final 5 min extension at the end of the reaction to ensure that all amplicons were completely extended and repeated for 40 amplification cycles. Relative quantification of the expression of relevant genes was performed using the ΔΔCt method after normalization to the expression of the housekeeping gene GAPDH. Primer sequences are listed in Supp Table 3.

***In vitro* assay:**

Cells were isolated from the MLN of control and cirrhotic rats, and FRC and T cells were sorted using FACS. FRCs were cultured in DMEM media supplemented with 10% FBS and 1% antibiotic solution, while T cells were maintained in suspension in RPMI media supplemented with 10% FBS and 1% antibiotic solution.

For co-culture experiments, cultured FRC were pretreated with recombinant versican (rVCAN) protein (RayBiotech, 230-00833-10) at a concentration of 2 ng/mL for 12 hrs. T cells (at a 1:10 FRC-to-T cell ratio) were added to control and rVCAN-treated FRCs and incubated for 6 hours. Following incubation, non-adherent T cells were gently removed by washing the coverslips three times with PBS. The adherent FRCs and T cells were then fixed with 4% PFA and stained. Imaging was performed using a Leica inverted confocal microscope. In a parallel experiment, T cells were washed, collected, and stained for T cell subsets, then analyzed by flow cytometry.

**T cell migration assay**

For T cell migration assays, transwell inserts were used. FRC were plated in the upper chamber of the transwell and allowed to adhere for 24 hours. Afterward, FRCs were treated with rVCAN for 6 to 12 hrs in serum-deprived media. T cells from control MLN were added to the upper chamber, while RPMI media supplemented with 10 ng/mL CCL21 or 5ng/mL of IL-6 was placed in the lower chamber. The co-cultures were incubated for 4 hours, and T cells in both the upper and lower wells were subsequently counted.

For CD44 receptor blockade, purified T cells were resuspended in complete RPMI-1640 medium supplemented with 10% FBS and incubated with a monoclonal anti-rat CD44 blocking antibody (V3S-0822-YC741, 5µg/1 million T cells) for 30 minutes at 4 °C. After incubation, cells were washed twice with PBS to remove unbound antibody and subsequently used for T cell migration assays. Isotype-matched IgG antibodies served as negative controls.

**T cell Suppression Assay**

Mononuclear cells were isolated from the MLN of control rats and stained with CD45, CD3, CD25, and CD134 antibodies. Activated T cells (CD3⁺CD134⁺) and Tregs (CD3⁺CD25⁺) were sorted by FACS and co-cultured for suppression assays. CD134⁺ T cells were first activated *in vitro* with PMA (10 ng/mL) and ionomycin (1 µg/mL) for 4hrs, and washed thoroughly, and then cultured with Tregs at a 1:1 ratio in the presence or absence of recombinant VCAN (rVCAN, 2 ng/mL).
A separate condition included CD134⁺ T cells cultured with rVCAN but without Tregs. After 24 hours, cells were harvested and stained for CD134 expression. The percentage of CD3⁺CD134⁺ T cells was quantified by flow cytometry. The degree of T cell suppression was calculated using the following equation:

S0 = %Activated CD134^+^ T cells

Sv = %Activated CD134^+^ T cells + VCAN (2ng/mL)

St = %Activated CD134^+^ T cells + %CD25^+^ Tregs

Stv = %Activated CD134^+^ T cells + %CD25^+^ Tregs + VCAN (2ng/mL)

%Suppression(+VCAN) =100×(1−Sv/Stv)

%Suppression(- VCAN)=100×(1−S0/St)

**Quantification of cytokine levels**

MLN samples were homogenized in RIPA lysis buffer, and total protein concentration was determined using the BCA Protein Estimation Kit (Thermo Fisher Scientific). For ELISA, 5 µg of total protein from each MLN lysate was added per well. Samples were diluted with the diluent provided in the kit, or with distilled water if no diluent was supplied.

Blood samples were collected in heparin-coated tubes and centrifuged at 3500 rpm to separate plasma, which was subsequently stored at -80 °C until analysis. Plasma concentrations of TNF-α (DY510-05R&D Systems), TGF-β (E01689Ra, BT Lab), IL-6 (DY506-05, R&D Systems), IL-10 (E0108Ra, BT Lab), and Versican (E2321Ra, BT Lab) were quantified using commercially available ELISA kits according to the manufacturers' protocols.

**Statistical Analysis**

Continuous variables are expressed as either mean±standard deviation for a continuous distribution or as median values for a skewed distribution. Continuous variables were compared between the two groups using an unpaired two-tailed Student's t-test or Mann-Whitney U test. Variables greater than two were compared using a one-way ANOVA followed by a post hoc Tukey test. Bar diagrams with various data points, dot plots, and box whisker plots were plotted using GraphPad Prism (version 8.0.1.GraphPad Software, San Diego, CA, USA), and statistical analysis was performed using GraphPad Prism. Statistical significance was set at p<0.05.

**Clinical Study**

**Patient Inclusion and Exclusion Criteria**

This prospective observational study was conducted on a cohort of 86 critically ill cirrhosis patients admitted to the HUD or ICU of the Institute of Liver and Biliary Sciences (ILBS), New Delhi, over a period of one year. All patients had decompensated liver disease, defined by the presence of ascites, hepatic encephalopathy, variceal bleeding, or jaundice. Adult patients (≥18 years) with histological, clinical, biochemical, and/or radiological confirmed cirrhosis were included. Patients were followed prospectively for 28 days or until the occurrence of in-hospital death, liver transplantation, or transjugular intrahepatic portosystemic shunt (TIPS). Participants with a previous history of liver transplantation (LT), hepatocellular carcinoma or other malignancies, active non-hepatic chronic inflammatory or autoimmune diseases, portal vein thrombosis, or who were pregnant, were excluded from the study. Informed written consent was obtained from all participants or their legally authorized representatives, and the study protocol was approved by the Institutional Ethics Committee of ILBS (IEC/2024/116/MA05).

**Study Design**

Demographic, clinical, and laboratory data were prospectively recorded at the time of admission for all enrolled patients. At baseline, detailed clinical assessment and standard laboratory investigations, including liver function tests, renal profile, coagulation parameters, and inflammatory markers, were performed. Plasma samples were collected on the day of admission and stored at -80°C. Patients were classified into acute decompensation (AD) and acute-on-chronic liver failure (ACLF) groups according to the CLIF-OF criteria. Organ failures were defined as follows: liver (serum bilirubin ≥12.0 mg/dl), renal (creatinine ≥2 mg/dl or renal replacement therapy), brain (HE III IV grade on West Heaven criteria), coagulation (INR ≥ 2.5 or platelet<20,000), circulatory (required vasopressor), and lung (PaO_2_/FiO_2_ ≤ 200). Based on these criteria, patients with failure of one or more organs were categorized as having ACLF, while those without organ failure but with acute decompensation events, including ascites, hepatic encephalopathy, gastrointestinal bleeding, AKI, or infection, were classified as AD. The severity of liver disease was assessed using CTP, MELD-Na, and CLIF-OF scores. All patients were monitored for 28 days or until death, liver transplantation, or TIPS, whichever occurred first, to evaluate short-term outcomes and predictors of mortality.

**Sample Collection and Laboratory Measurements**

Blood samples were collected on the day of admission and centrifuged at 2,000 rpm for 10 minutes; the supernatant was stored at -80°C until further analysis. VCAN levels were quantified using a commercially available Human VCAN ELISA kit (BT Lab, E1909Hu).

**Statistical Analysis**

Categorical variables are presented as counts and percentages, and continuous variables as mean (25th–75th percentiles) unless otherwise specified. Between-group comparisons were performed using Pearson's chi-square test for categorical variables, and Student's t-test or the Mann-Whitney U test for continuous variables for two groups and by one-way Anova or Kruskal-Wallis test for more than 2 groups, depending on the distribution. Differences in VCAN levels between survivors and non-survivors and among clinical outcome groups were evaluated using the Mann-Whitney U test. Survivors were defined as patients alive or discharged within **28 days** of enrollment. Clinical predictors were entered into the Cox hazard regression model to assess the effects of factors on 28-day in-hospital mortality. We used unadjusted linear regression models and multivariate models adjusted for clinical covariates. In univariate analysis, factors with a significance of p<0.2 were entered into multivariate analysis. Results are reported as HR with 95% CIs and p-values <0.5 considered significant. ROC curves were constructed to evaluate the discrimination of VCAN and other candidate biomarkers; the AUC with 95% CIs was reported. Optimal cut-off values were selected using the Youden index (maximum [sensitivity + specificity - 1]) and reported with corresponding sensitivity and specificity. For the visualization of time-to-event data, Kaplan-Meier curves were used for unadjusted survival comparisons. However, because liver transplantation and TIPS placement can preclude the event of interest (death), competing-risk methods were employed for primary prognostic analyses.

Competing risk regression analysis was performed to evaluate the association between VCAN levels and mortality, accounting for competing events (TIPS or transplantation). The Fine and Gray proportional subdistribution hazards model was used to estimate the subdistribution hazard ratio (SHR) for the cumulative incidence of the primary event (i.e., mortality), treating competing outcomes as potential risks rather than censoring events. VCAN levels were analyzed as a continuous variable, and univariate analysis was initially performed to identify predictors associated with outcome events. Variables with statistical significance (p < 0.2) in univariate analysis were included in a multivariate Fine-Gray model to adjust for potential confounders. Regression coefficients, standard errors, Z-statistics, p-values, and 95% CIs were computed. The SHR ratio represented the relative change in hazard per unit increase in VCAN concentration. Covariate evaluation points were calculated for the 25^th^, 50^th^, and 75^th^ percentiles of VCAN level (14.874, 17.370, and 19.060, respectively). Cumulative incidence functions were plotted for these quantiles to display the estimated probability of the outcome over time while accounting for the presence of competing risks. All analyses were conducted in Stata (version 17; StataCorp LLC) using the Fine and Gray model and cumulative incidence estimation for competing risks regression, implemented in R (CRR function within the cmprsk package).

**Supplementary Figure Legends**

**Supplementary Figure 1. Bacterial load in the rats increased after MLN resection.** (a) Preparation and characterization of an animal model of liver cirrhosis. CCl4:olive oil in a ratio of 1:1 was injected intraperitoneally in 6-week-old SD rats for 12 weeks. At the end of 12 weeks, 1mg/kg LPS was injected i.p., and rats were sacrificed 24 h post-LPS. (b) HE and MT staining of liver from control, C+LPS, and CCl4+LPS rats. (c, d) Detection and quantification of endogenous bacterial load in MLN and extraintestinal organs, portal blood, and efferent mesenteric lymph in healthy control and 12-week CCl₄-induced cirrhotic rats. (e) Schematic showing MLN resection in control rats (CMLNx) followed by acute LPS challenge and sample collection 24h post LPS. (f-g) Detection and quantification of endogenous bacterial load in 100 mg tissue of liver, spleen, and lungs of C+LPS and CMLNx+LPS rats. (h) MT and HE staining of the liver section of the CCl₄ and CCl₄-MLNx rat. (i) Quantification of collagen deposition in liver sections of CCl4 and CCl-MLNx rats (j) Kaplan–Meier survival curve of CCl₄-MLNx and CCl_4_ rats along with at risk table. Data pooled from 4 independent experiments and represented as mean±SD. Statistical comparisons were made using the (g) one-way ANOVA with post-hoc Tukey's multiple comparison test, (i) unpaired two-tailed Student's *t*-test, and (j) log-rank test. *p < 0.05, **p < 0.01, and ***p < 0.001. MLN: Mesenteric lymph node.

**Supplementary Figure 2: Gating strategy for quantification of immune cells from MLN, spleen, blood, and lymph from experimental groups.** For quantification of (a) T cells, (b) monocytes and dendritic cells, and (c) macrophages.

**Supplementary Figure 3. Quantification of immune cell subsets in MLN, systemic circulation, lymphatic and portal compartments following LPS stimulation in control and cirrhotic rats.** (a, b) Representative plots and quantification of CD4+ and CD8+ T cell subsets in the (a) MLN and (b) peripheral blood. (c-f) Representative plots and quantification of (c) DC, (d) monocytes, and (e-f) T cells in portal blood of control, C+LPS, and CCl4+LPS groups. (g-h) Representative plots and quantification of T cells in the efferent mesenteric lymph of the studied groups**.** Data pooled from ≥3 independent experiments and expressed as mean±SD. Statistics were accessed by using (c) one-way ANOVA, (a, b, and d-h ) two-way ANOVA with post-hoc Tukey's test. ***p < 0.05, ****p < 0.01, *****p < 0.001.** * vs control; # vs C+LPS.

**Supplementary Figure 4. Immune profiling in systemic and portal circulation following MLN resection and LPS administration.** (a) Representative plots and quantification of CD4+ and CD8+ T cell subsets in the peripheral blood of C+LPS and CMLNx+LPS rats. (b-e) Representative plots and quantification of (b) DC, (c) monocytes, and (d-e) T cells in portal blood of C+LPS and CMLNx+LPS rats. (f) Representative plots and quantification of CD4+ and CD8+ T cell subsets in the portal blood of CCl4+LPS and CCl4-MLNx+LPS rats. (g-j) Representative plots and quantification of (g) DC, (h) monocytes, and (i-j) T cells in portal blood of CCl4+LPS and CCl4-MLNx+LPS rats. (k-n)Immune cell quantification in the spleen of all studied groups. Representative plots show quantification of (k) CD103+CD80+DC, (l) CD43+ monocytes, (m) CD68+macrophages, (n) CD4+ and CD8+T cells. (o) mRNA expression of cytokines in spleen of C+LPS vs CMLNx+LPS and CCl4+LPS vs CCl4-MLNx+LPS groups. Data pooled from ≥3 independent experiments and expressed as mean±SD. Statistics were accessed by using (b, g, o) unpaired two-tailed Student's *t*-test, (k, m) one-way ANOVA, (a, c-e, f, h-j, l, n ) two-way ANOVA with post-hoc Tukey's test. ***p < 0.05, ****p < 0.01, *****p < 0.001.** DC: Dendritic cells. MLNx: MLN resected rats; MLN: Mesenteric lymph node.

**Supplementary Figure 5. Molecular proteomic profiling and cellular source of VCAN in MLN of control and cirrhotic rats.** (a) PCA plot displaying distinct clustering of MLN samples from control, C+LPS, and CCl₄+LPS groups (n=2 each) based on global protein expression profiles, indicating group-specific molecular signatures. Proteomics was done from 3 MLN samples, each group, of which 2 were selected from each group for further analysis. (b) Pathway enrichment analysis showing fold change values for selected proteins involved in immune regulation, chemotaxis, cytoskeletal remodeling, and negative regulation of cellular migration, upregulated in CCl₄+LPS vs C+LPS MLN. Differentially expressed proteins were identified using a fold change threshold ≥1.5 and adjusted p-value (P_adj_) < 0.2. (c) Quantification of fibrosis in the liver and MLN of BDL models at different time points via ImageJ (one point in the graph represents the mean of the %area quantified from 3 different sides of one section of tissue). (d) VCAN levels in MLN lysate of BDL rats with and without bacterial load in MLN, n=6 each. Data expressed as mean±SD. Statistics were accessed by using (c) one-way ANOVA with post-hoc Tukey's test, and (d) unpaired two-tailed Student's *t*-test. ***p < 0.05, ****p < 0.01, *****p < 0.001.** MLN: Mesenteric lymph node.

**Supplementary Figure 6. Immune profiling in MLN, systemic, and spleen after VCAN knockdown using siRNA and LPS administration.** (a) Schematic representation of the experimental workflow for isolation and sorting of MLN cells into immune cells, endothelial cells, and stromal FRC to identify the source of VCAN expression. (b) mRNA expression of VCAN in sorted immune, endothelial, and FRC subsets from MLN of a control rat. (c) mRNA expression of VCAN in sorted FRC subsets from control, C+LPS, and CCl4+LPS rats. (d) Relative gene expression of VCAN in MLN of ctrl-siRNA and VCAN-siRNA treated rats. (f) Quantification of DC, monocytes, macrophages, and T cells in MLN of ctrl-siRNA and VCAN-siRNA treated rats. (g) Cytokine level in MLN lysate of ctrl-siRNA and VCAN-siRNA treated rats. (h) Quantification of DC, monocytes, and T cells in the peripheral blood of ctrl-siRNA and VCAN-siRNA treated rats. (i) Quantification of DC, monocytes, macrophages, and T cells in the spleen of ctrl-siRNA and VCAN-siRNA treated rats. (j) Cytokine level in plasma of ctrl-siRNA and VCAN-siRNA treated rats. Data pooled from ≥3 independent experiments and expressed as mean±SD. Statistics were accessed by using (d and f-j) unpaired two-tailed Student's *t*-test, (b, c) one-way ANOVA, (f, h, and i) two-way ANOVA with post-hoc Tukey's test. ***p < 0.05, ****p < 0.01, *****p < 0.001**. FRC: fibroblast reticular cells; MLN: Mesenteric lymph node; DC: Dendritic cells.

**Supplementary Tables:**

**Supp Table 1: DEP in proteomic analysis of MLN from Control, C+LPS and CCl_4_+LPS rats:**

| **Cvs** CCl_4_**+LPS upregulated and C+LPSvsCCl4+LPS upregulated** | | | | | | | | |
| --- | --- | --- | --- | --- | --- | --- | --- | --- |
| **Accession Numbers** | Protein_grpID | Control 1_intensity | Control 3_intensity | C+LPS1  intensity | C+LPS2  intensity | CCl4+LPS2  intensity | CCl4+LPS3  intensity | Description |
| A0A0G2JTM0 | 1134 | 21.9 | 10.2 | 1.5 | 4.5 | 624.2 | 665 | Acid-sensing ion channel 2 OS=Rattus norvegicus OX=10116 GN=Asic2 PE=4 SV=1 |
| A0A0G2JZT1 | 253 | 81.3 | 72 | 61.7 | 59.6 | 121.5 | 108.4 | NACHT and WD repeat domain-containing 2 OS=Rattus norvegicus OX=10116 GN=Nwd2 PE=4 SV=2 |
| A0A0G2K7Z1 | 11839 | 14.9 | 9.4 | 13.2 | 12.4 | 447.4 | 521.7 | Zinc finger protein 346 OS=Rattus norvegicus OX=10116 GN=Zfp346 PE=1 SV=1 |
| A0A8I5ZTJ6 | 2339 | 35 | 24.3 | 25.4 | 44.7 | 1050.3 | 906.4 | Casein kinase 2 subunit alpha'-interacting protein OS=Rattus norvegicus OX=10116 GN=Csnka2ip PE=4 SV=1 |
| B0BN44 | 5552 | 41.8 | 23.4 | 18.1 | 9.3 | 141.1 | 151.8 | Tumor protein p53-inducible protein 13 OS=Rattus norvegicus OX=10116 GN=Tp53i13 PE=2 SV=1 |
| D3ZJH1 | 15536 | 6.7 | 22.3 | 23.3 | 27 | 413.2 | 422.1 | C-C motif chemokine receptor 10 OS=Rattus norvegicus OX=10116 GN=Ccr10 PE=3 SV=1 |
| D3ZPZ3 | 18675 | 30.3 | 14.6 | 39.5 | 42.2 | 1198.6 | 1403.6 | DNA-(apurinic or apyrimidinic site) endonuclease OS=Rattus norvegicus OX=10116 GN=Apex2 PE=3 SV=4 |
| D4A8V2 | 9610 | 12.8 | 24.4 | 13.4 | 27.5 | 290.4 | 320.8 | Coiled-coil domain-containing 177 OS=Rattus norvegicus OX=10116 GN=Ccdc177 PE=1 SV=1 |
| Q06605 | 9325 | 171 | 81.7 | 57.8 | 95.6 | 371 | 437.1 | Granzyme-like protein 1 OS=Rattus norvegicus OX=10116 PE=2 SV=1 |
| Q3MIE0 | 17280 | 6.3 | 10.2 | 10.8 | 7.9 | 680.7 | 859.3 | Enoyl-CoA hydratase domain-containing protein 3, mitochondrial OS=Rattus norvegicus OX=10116 GN=Echdc3 PE=2 SV=1 |
| Q5XI94 | 8309 | 8.9 | 3.7 | 38.6 | 22.7 | 83.2 | 93.1 | Uncharacterized protein C1orf112 homolog OS=Rattus norvegicus OX=10116 PE=2 SV=1 |
| Q62769 | 19444 | 22.2 | 18.9 | 21.5 | 20.4 | 1110.7 | 1413.7 | Protein unc-13 homolog B OS=Rattus norvegicus OX=10116 GN=Unc13b PE=1 SV=2 |
| Q641Z2 | 18877 | 32.2 | 25.4 | 161.2 | 69 | 447.4 | 522 | Tyrosine-protein phosphatase non-receptor type 9 OS=Rattus norvegicus OX=10116 GN=Ptpn9 PE=2 SV=1 |
| Q6AYU0 | 19390 | 107.5 | 92.7 | 21.8 | 44.3 | 241 | 333.5 | Uncharacterized protein C12orf50 homolog OS=Rattus norvegicus OX=10116 PE=2 SV=1 |
| Q9ERB4 | 14789 | 67 | 62 | 61.3 | 22.4 | 405.2 | 529.6 | Versican core protein (Fragment) OS=Rattus norvegicus OX=10116 GN=Vcan PE=2 SV=2 |
| Q9JHU0 | 5480 | 1.4 | 1.1 | 4.9 | 2.7 | 45.5 | 33.2 | Dihydropyrimidinase-related protein 5 OS=Rattus norvegicus OX=10116 GN=Dpysl5 PE=1 SV=1 |
| **CvsC+LPS upregulated and C+LPSvsCCl4+LPS downregulated** | | | | | | | | |
| A0A8I6AJM0 | 4968 | 3.1 | 2.9 | 139.8 | 150.9 | 7.9 | 12.4 | G protein-regulated inducer of neurite outgrowth 1 OS=Rattus norvegicus OX=10116 GN=Gprin1 PE=4 SV=1 |
| A2RUW0 | 5349 | 112.2 | 150.9 | 357 | 326.3 | 141.7 | 54.9 | Maestro heat-like repeat-containing protein family member 7 OS=Rattus norvegicus OX=10116 GN=Mroh7 PE=1 SV=1 |
| B2GV05 | 13653 | 1.5 | 2 | 28.8 | 32.5 | 1.5 | 1.9 | RNA-binding protein 5 OS=Rattus norvegicus OX=10116 GN=Rbm5 PE=1 SV=1 |
| B2RYG7 | 18449 | 115.7 | 110.4 | 240.9 | 275.5 | 25.4 | 66.7 | Docking protein 3 OS=Rattus norvegicus OX=10116 GN=Dok3 PE=1 SV=1 |
| B5DEL3 | 18896 | 20.2 | 12.8 | 313.4 | 382.2 | 15.4 | 15.3 | Tetratricopeptide repeat protein 17 OS=Rattus norvegicus OX=10116 GN=Ttc17 PE=1 SV=1 |
| D3ZHU3 | 8138 | 99.3 | 127 | 358.7 | 390.3 | 129.9 | 78.2 | MEF2-activating motif and SAP domain-containing transcriptional regulator OS=Rattus norvegicus OX=10116 GN=Mamstr PE=4 SV=1 |
| D3ZR46 | 14241 | 3.5 | 11.9 | 44.6 | 44 | 14.6 | 12.8 | Serine protease 40 OS=Rattus norvegicus OX=10116 GN=Prss40 PE=4 SV=1 |
| D4AD22 | 11972 | 11.7 | 12.6 | 288.3 | 361.7 | 14.3 | 13.5 | Otolin 1 OS=Rattus norvegicus OX=10116 GN=Otol1 PE=4 SV=1 |
| E9PU17 | 5880 | 117.4 | 87.6 | 197.8 | 193.5 | 61.5 | 96.4 | ATP-binding cassette sub-family A member 17 OS=Rattus norvegicus OX=10116 GN=Abca17 PE=2 SV=1 |
| P35289 | 17706 | 0.7 | 2.2 | 34 | 46.3 | 5.2 | 3.7 | Ras-related protein Rab-15 OS=Rattus norvegicus OX=10116 GN=Rab15 PE=2 SV=1 |
| P60881 | 9545 | 5.2 | 12.4 | 31.9 | 34.4 | 7.7 | 2.3 | Synaptosomal-associated protein 25 OS=Rattus norvegicus OX=10116 GN=Snap25 PE=1 SV=1 |
| Q32ZH1 | 10752 | 18.1 | 6.2 | 122.7 | 146.8 | 7.1 | 14 | Beta-defensin OS=Rattus norvegicus OX=10116 GN=Defb21 PE=2 SV=1 |
| Q4KSH7 | 8304 | 192.9 | 130.2 | 339.8 | 328.8 | 47.6 | 95.4 | Dual specificity mitogen-activated protein kinase kinase 7 OS=Rattus norvegicus OX=10116 GN=Map2k7 PE=1 SV=1 |
| Q99P84 | 1811 | 211.5 | 224.6 | 421.1 | 434.9 | 269.6 | 188.7 | 1-phosphatidylinositol 4,5-bisphosphate phosphodiesterase epsilon-1 OS=Rattus norvegicus OX=10116 GN=Plce1 PE=1 SV=1 |
| Q9EPU2 | 3621 | 17.9 | 22 | 1167.9 | 1138.3 | 14.8 | 19.7 | Nuclear receptor coactivator 3 (Fragment) OS=Rattus norvegicus OX=10116 GN=Ncoa3 PE=2 SV=1 |
| Q9JI66 | 9065 | 7.7 | 7.7 | 551.7 | 486 | 7.1 | 6.1 | Electrogenic sodium bicarbonate cotransporter 1 OS=Rattus norvegicus OX=10116 GN=Slc4a4 PE=1 SV=1 |
| **CvsCCl4+LPS downregulated and C+LPSvsCCl4+LPS downregulated** | | | | | | | | |
| D3ZSZ3 | 19440 | 144 | 155.4 | 121.6 | 134.3 | 56.7 | 31.6 | Serine/threonine-protein kinase NLK OS=Rattus norvegicus OX=10116 GN=Nlk PE=3 SV=1 |
| D3ZZN3 | 17523 | 34.3 | 45.4 | 49.6 | 56.5 | 7.5 | 7.9 | Acetyl-coenzyme A synthetase OS=Rattus norvegicus OX=10116 GN=Acss1 PE=3 SV=1 |
| Q5BKC3 | 5438 | 383.3 | 320.5 | 361.8 | 358.7 | 7.8 | 10.1 | Protein deglycase OS=Rattus norvegicus OX=10116 GN=Park7 PE=1 SV=1 |
| Q5FVF9 | 18630 | 304.6 | 310.7 | 432.4 | 382.4 | 204.1 | 122.7 | Biotinidase OS=Rattus norvegicus OX=10116 GN=Btd PE=2 SV=1 |
| Q64305 | 7456 | 63.9 | 63.7 | 49.3 | 59.3 | 3 | 3.3 | Pancreas transcription factor 1 subunit alpha OS=Rattus norvegicus OX=10116 GN=Ptf1a PE=1 SV=1 |

**Supp Table 2: Antibody Panel**

| **Antibody** | **Brand** | **Catalogue** | **Fluorochrome** |
| --- | --- | --- | --- |
| Podoplanin | BioLegend | 250906 | APC |
| CD31 | BD Biosciences | 555027 | PE |
| CD45 | BioLegend | 202216 | APC-Cy7 |
| CD3 | BioLegend | 201413 | APC |
| CD4 | BioLegend | 201519 | PE-Cy7 |
| CD8 | BioLegend | 201715 | PerCP-Cy5.5 |
| CD25 | BioLegend | 202103 | PE |
| CD134 | BioLegend | 204508 | FITC |
| CD11b | BioLegend | 201817 | PE-Cy7 |
| CD103 | BioLegend | 205509 | APC |
| CD80 | BioLegend | 200205 | PE |
| CD68 | BioLegend | 201003 | PE |
| CD44 | Creative Biolabs | V3S-0822-YC741 | Used for Blocking |

**Supp Table 3: List of rat genes and primers used for qRT-PCR**

| **Gene** | **Forward Primer** | **Reverse Primer** |
| --- | --- | --- |
| **IL-6** | TTGTTGACAGCCACTGCCTTC | GAATTGCCATTGCACAACTCTTTTC |
| **IL-10** | TAAGGGTTACTTGGGTTGCC | TATCCAGAGGGTCTTCAGC |
| **TNF-α** | AAATGGGCTCCCTCTCATCAGTTC | TCTGCTTGGTGGTTTGCTACGAC |
| **TGF-β** | GCTAATGGTGGACCGCAACAAC | TGGCACTGCTTCCCGAATGTC |
| **VCAN** | ACGAATACCCTCGCAGAAAC | TCGGTGACTAATGGAATGACTG |
| **S1PR1** | TTCAGCCTCCTTGCTATCGC | AGGATGAGGGAGATGACCCAG |
